# The mosquito microbiome includes habitat-specific but rare symbionts

**DOI:** 10.1101/2021.09.24.461628

**Authors:** Hans Schrieke, Loïs Maignien, Florentin Constancias, Florian Trigodet, Sarah Chakloute, Ignace Rakotoarivony, Albane Marie, Gregory L’Ambert, Patrick Makoundou, Nonito Pages, A. Murat Eren, Mylène Weill, Mathieu Sicard, Julie Reveillaud

## Abstract

Microbial communities are known to influence mosquito lifestyles by modifying essential metabolic and behavioral processes that affect reproduction, development, immunity, digestion, egg survival, and ability to transmit pathogens. Many studies have used 16S rRNA gene amplicons to characterize mosquito microbiota and investigate factors that influence host-microbiota dynamics. However, a relatively low taxonomic resolution due to clustering methods based on arbitrary threshold and the overall dominance of *Wolbachia or Asaia* populations obscured the investigation of rare members of mosquito microbiota in previous studies. Here, we used high resolution Shannon entropy-based oligotyping approaches to analyze the microbiota of *Culex pipiens, Culex quinquefasciatus* and *Aedes* individuals from continental and overseas regions in Southern France and Guadeloupe as well as from laboratories with or without antibiotics treatment. Our experimental design that resulted in a series of mosquito samples with a gradient of *Wolbachia* density and relative abundance along with high-resolution analyses of amplicon sequences enabled the recovery of a robust signal from typically less accessible bacterial taxa. Our data confirm species-specific mosquito-bacteria associations with geography as a primary factor that influences bacterial community structure. But interestingly, they also reveal co-occurring symbiotic bacterial variants within single individuals for both *Elizabethkingia* and *Erwinia* genera, distinct and specific *Asaia* and *Chryseobacterium* in continental and overseas territories and a putative rare *Wolbachia* variant. Overall, our study reveals the presence of previously-overlooked microdiversity and multiple closely related symbiotic strains within mosquito individuals with a remarkable habitat-specificity.

## INTRODUCTION

In our changing world, global warming, habitat reduction and increase of transport impact host-parasite interactions and support a global emergence of wildlife zoonotic diseases^1–3^. In this context, mosquitoes represent a significant threat to global public health because of their ability to transmit multiple pathogens including eucaryotes, bacteria and viruses. Among the most well-known arthropod-borne viruses or arboviruses, Dengue, West Nile, Zika and chikungunya are responsible for several hundreds of thousands of deaths and billions of cases every year^4^. Because of a widespread insecticide resistance^5,6^ and in the absence of efficient vaccines or therapy for now, manipulation of the microbiota, and in particular of mosquito bacterial symbionts are emerging as novel vector control strategies^7–10^.

The maternally inherited endosymbiotic *Wolbachia*, which is able to modify the host reproduction and limit the transmission of pathogens, can be an effective biological weapon against arboviruses. Introduction of infected individuals in local population is a cornerstone for all the techniques using *Wolbachia* either to block the pathogens or sterilizing the females^11^. It is therefore critical to understand the evolution of *Wolbachia* in natural population before deploying these methods at larger scales and believing in their sustainability. For this, fine-tuned studies of *Wolbachia* population diversity variants are required. Likewise, other bacterial symbionts can interfere with mosquito-pathogen interactions and offer great potential for paratransgenic applications, *i*.*e*, the use of engineered symbionts to express anti-pathogenic molecules^12^. For example, *Asaia* sp. can activate the expression of antimicrobial peptides in *Anopheles stephensi* post *Plasmodium* infection^13^. On the one hand, the intestinal bacterium *Serratia marcescens* enhances the vectorial susceptibility to arbovirus in *Aedes* mosquitoes by secreting the SmEnhancin protein that damages the physical barrier to the Dengue viruses, therefore facilitating the infection and spread of the virus^14^. On the other hand, the strain Y1 from the same intestinal bacterium can inhibit the development of *Plasmodium* by activation of the immune system in *Anopheles stephensi* mosquitoes^15^. These studies highlight the importance of a resolution at the strain level during microbiota analysis.

In addition, some of these mosquito bacterial symbionts and other microbial communities also have key roles on the development and physiology of mosquito species. Bacterial communities have major functions in mosquito growth and larval development^16,17^, blood and sugar digestion^18^, immunity^19,20^ and egg production^18^. Therefore, a comprehensive understanding of the diversity, potential diverse functions and possible interactions between the different microbiota members, as well as between the microbiota, pathogens and their hosts is key. Critical to the manipulation of mosquito microbiota as a novel vector control approach is an *in-depth* knowledge of their microbiota.

However, mosquito samples are often dominated by a symbiont, that is, *Wolbachia* or *Asaia* in *Culex* and *Aedes*, respectively, hampering a holistic characterization of the microbiota and notably of the less abundant but possibly important rare taxa. The rare members of the microbiota are also important reservoir of genetic and functional diversity with key ecological roles, as shown in the marine and the terrestrial environment as well as in the plant and the human microbiome^21–23^. Because most studies of high-throughput sequencing marker genes datasets use clustering methods, often based on an arbitrary threshold of 97 to 98%, they can overlook closely related ecologically relevant sequence variants. The recently developed Minimum Entropy Decomposition (MED) and oligotyping methods use the Shannon entropy to identify ecologically relevant units at the whole community or at a taxa level (from phylum to species level), in an unsupervised and supervised manner, respectively^24,25^. These information-theory based approaches have allowed unveiling previously undetected ecological patterns for microbial communities from diverse free-living and host-associated environments including sewages, the sponge and coral microbiome, as well as the oral and plaque microbiota among others^24,26–28^. Recently, Coon and colleagues^29^ suggested bacterial communities differed substantially in *Aedes* and *Culex* larvae from different collection sites in the United States using oligotyping, although they focused on pools of larvae. Overall, the single-nucleotide resolution of MED and oligotypes make it possible to reveal novel bacterial variants, with high level of host specificity and to characterize bacterial populations in a microdiversity-aware manner.

To gain deeper insights into typically overlooked rare microbial symbionts in mosquitoes, here we used a high-resolution sampling and analysis strategy to study their microbial community compositions. We investigated microbial communities of *Aedes aegypti, Culex pipiens* and *Culex quinquefasciatus* whole mosquito individuals and their reproductive organs from field (continental and overseas regions of Southern France and Guadeloupe) and insectaries (including some that were antibiotically-treated) to obtain a gradient of *Wolbachia* relative abundance. Together with fine scale entropy-based MED and oligotyping approaches as well as metagenome screening, we aimed to get access to the less abundant yet possibly functionally important mosquito-associated bacterial taxa.

## MATERIALS AND METHODS

### Specimen collection and dissection

We collected field mosquito individuals using a carbon dioxide mosquito trap in Languedoc, Herault, France (Bosc Viel in Mauguio and Camping l’Europe Vic La Gardiole, with the help of Entente Interdépartementale de Démoustication Méditerranée EID) and in Guadeloupe (Prise d’eau, Commune de Petit Bourg) in 2017 and 2018. We transported specimens alive to the laboratory immediately upon recovery. Laboratory specimen *Culex pipiens* from Lavar (St Etienne, France) and *Culex quinquefasciatus* (Slab treated by Tetracycline at five generations before our experiments, originally from Southern California, also called Slab TC) were reared at Institute of Evolutionary Science of Montpellier insectarium (ISEM, Montpellier, France).

We afforded mosquito manipulation and dissection as described in^30^. Briefly, we anesthetized adult females for 4 min at −20 °C and surface-sterilized insects through gentle vortexing with cold (4 °C) 96% ethanol. We transferred them into a sterile cold (4 °C) phosphate-buffered saline (PBS) 1× solution and then onto a sterile microscope slide with sterile PBS 1× on top of a cold plate prior to dissection of ovaries using sterilized tweezers. We stored whole or dissected mosquito specimen at −80 °C until further processing.

### DNA extraction, V3-V4 polymerase chain reaction (PCR) and Illumina tag sequencing

We extracted total genomic DNA in a sterile hood with the DNeasy Blood and Tissue (Qiagen) following manufacturer instructions. We systematically added one extraction blank (Negative Control, hereafter blank), corresponding to an extraction without sample tissue, to each of a series of 10 DNA extraction microtubes. We quantified DNA with a QUBIT 2.0 Fluorometer and diluted overconcentrated samples to attempt generating amplicon librairies from equimolar DNA solutions for each run (pools of either 1ng.ul or 30 ng.ul). We conducted all pre-PCR laboratory manipulations with filtered tips under a sterile environment in a DNA-free room dedicated to the preparation of PCR mix and equipped with hoods that are kept free of DNA by UV irradiation. We used primers and the dual-index method^31^ to amplify a 429-bp portion of the V3-V4 region of the 16S rRNA gene. The combination of primers including different 8-bp indices (8 i5-indexed forward primers and 12 i7-indexed reverse primers) allowed multiplexing 96 samples onto three MiSeq flow cells.

We analyzed a total of 136 samples including 71 whole specimens and 50 dissected ovaries (where vertically transmitted symbionts can be found) together with 15 blanks (See Supplementary Table 1). Whole specimen samples consisted of 43 *Culex pipiens* (7 individuals from Camping Europe, 14 from Bosc and 22 from Lavar), 18 *Culex quinquefasciatus* samples (4 from Guadeloupe, and 14 Slab TC) and 10 *Aedes aegypti* from Guadeloupe. Among the ovaries, we counted 44 *Culex pipiens* samples (21 from Camping Europe, 10 from Bosc and 13 from Lavar), 3 *Culex quinquefasciatus* and 3 *Aedes aegypti* samples, both from Guadeloupe).

### Identification of oligotypes, filtration of contaminants, and taxonomic assignment

We merged and quality-filtered raw sequences using the Illumina-Utils pipeline version 2.8^32^ with stringent settings (min-overlap-size 30 --max-num-mismatches 0 --enforce-Q30). We used the algorithm minimum entropy decomposition (MED)^24^ with default parameters to identify sequence variants in high-throughput sequencing of 16S rRNA gene amplicons. MED is an algorithm to partition a given set of sequences into discrete sequence groups (terminal MED nodes, hereafter unsupervised oligotypes (UO)) by minimizing the total entropy in the dataset^24^. We identified and removed contaminant UO that could come from several sources (laboratory manipulation, DNA extraction kit or PCR reagents kitomes^33^) using the prevalence method from the R package Decontam version 1.6.0^34^. We computed a statistical score based on a presence / absence table of UO counts in true samples and blanks. We used a threshold of 0,65 to remove contaminants, which consisted of UO with higher prevalence in blanks than in samples (Supplementary Figure 1). We assigned taxonomy to the representative sequence of each retained UO using the *assignTaxonomy()* function (minBoot=80) based on the RDP Naive Bayesian Classifier algorithm^35^ and the *addSpecies()* function from the R package Dada2 version 1.14.1^36^ using the silva138 reference database^37^.

In cases where maximum sensitivity was essential, we further investigated genus-level UO generated from our data by manually identifying nucleotide positions of high entropy using the supervised oligotyping method^25^, which enables supervised input into the curation of final results. Briefly, we concatenated the sequences from all unsupervised oligotypes assigned to a specific genus and ran the oligotyping pipeline. Supervised oligotypes are defined as the concatenation of the high-information nucleotide positions identified by Shannon entropy with a threshold value of minimum 0.2. For this, we used the default component (-c) to identify the nucleotide positions with the highest Shannon entropy. We set the minimum substantive abundance threshold for a supervised oligotype (-M) to 10. We further confirmed these results for M=100 (data not shown). Random sequencing errors are reported to generate entropy values lower than 0.2 for Illumina data (https://merenlab.org/2013/11/04/oligotyping-best-practices/).

### Statistical analyses

We imported raw unsupervised oligotype count table, taxonomic assignments and metadata in the phyloseq package for data handling^38^ using R version 3.6^39^. We analyzed microbial communities using alpha diversity metrics including UO observed richness, Chao1’s estimated richness and Shannon’s diversity index on unnormalized count data. Both Observed and Chao1 indexes account for the number of species (richness) with the latter giving with more weight to rare species using a singletons to doubletons ratio while Shannon accounts for both abundance and evenness. Community similarity (i.e., beta-diversity) was quantified using Bray-Curtis distance on normalized count table (relative abundance) as in^40^ and visualized using NMDS ordination. Samples with a sequencing depth lower than 1000 reads were discarded (13/136) to allow an exhaustive description of bacterial communities with some exceptions (see Supplementary Note 1 for details). We tested differences in alpha and beta-diversity between species or location using ANOVA (Analysis of variance, *aov()* function from the basic package stats from R version 3.6^39^) and PERMANOVA statistical tests (*adonis()* function from the vegan R package version 2.5-6^41^), respectively. In order to test if sequencing depth influenced differences in alpha-diversity between groups, the latter was added in the model (alpha-diversity ∼ Group * Sequencing depth). A significant interaction between Group and Sequencing depth was detected only for the Shannon index (see Supplementary Table 2 - Sheet 1) indicating that sequencing depth influenced Shannon values among groups. Therefore, statistical tests for the Shannon index were performed on the rarefied dataset as well (using the *rarefy_even_depth*() function from the phyloseq package; 100 reads). Multiple group comparisons were performed using Tukey’s test (*glht()* function from multcomp R package version 1.4^42^) for alpha diversity and pairwise PERMANOVA for beta-diversity. We then generated rarefaction curves using the ggrare() function from the R package ranacapa^43^ to investigate sequencing depth efforts per sample.

### *Wolbachia* relative abundance and density in *Culex* specimen

We investigated the *Wolbachia* relative abundance from the high-throughput sequencing *Culex* dataset using a Hierarchical Cluster Analysis (or HCA) that limits the arbitrary choice of a threshold value to split data. For that, we used three functions from the basic stats package from R 3.6^39^: *dist()* to convert the percentage of *Wolbachia* per sample into an Euclidean distance matrix, *hclust()* to perform the HCA using the “ward.D2” method and *cutree()* to cut the tree resulting from the HCA into three groups (k=3, hereafter referred to “Low infection”, “Medium infection” and “High infection” groups. Using this approach, we ended up splitting samples with *Wolbachia* relative abundance between 0 and 5%, 10 to 69% and 69 to 100%, respectively (Supplementary Figure 2 and Supplementary Table 3 – Sheet 1).

In addition, we used real-time quantitative PCR (qPCR) on a subset of representative samples (n=34) in order to estimate the *Wolbachia* density and investigate its correspondence between the *Wolbachia* infection groups obtained with HCA. Briefly, we used *Wolbachia* wsp (using wolpipdir and wolpiprev primers as in^44^) and Culex Ace-2 genes (using acequantidir and acequantirev as in^45^) to estimate the normalized *Wolbachia* amount per Ace-2 (host) copy. Samples with a Ct (cycle threshold) value >33 were excluded to avoid uncertain low titer quantification.

### Screening ovary shotgun metagenomes for the presence of potentially distinct *Wolbachia* variants

To further validate the occurrence of putative distinct *Wolbachia* variants within the reproductive organs, we screened available ovary shotgun metagenomes with the identified supervised oligotypes. We extracted fragments of 30bp from the supervised oligotype sequences including the positions with highest entropy that were used to discriminate the different *Wolbachia* variants, and used the anvi’o^46^ program “anvi-script-get-primer-matches” to search for metagenomic sequences that include these fragments in four ovaries metagenomes of *Culex pipiens* mosquitoes from Southern France^30^ (European Nucleotide Archive ENA accession numbers: ERS2407346, Culex O03; ERS2407347, Culex O07; ERS2407348, Culex O11; ERS2407349, Culex O12) as well as egg-rafts from three *C. pipiens* isofemale lines from North Africa^47^ (SRR5810516, Pipiens_MGx_Istanbul; SRR5810518, Pipiens_MGx_Tunis; SRR5810517 Pipiens_MGx_Harash). We used the quality filtered versions of these datasets for more accuracy.

## RESULTS

### Estimated bacterial diversity

We obtained a total of 6,452,623 high-quality reads distributed into 67 unsupervised oligotypes from 113 samples, including 67 whole specimens (3,402,384 reads) and 46 ovary samples (3,050,239 reads) after stringent quality filtering of Illumina reads and decontamination (see Supplementary Note 1 for the detailed bioinformatic analyses). Supplementary Figure 3 and Supplementary Table 4 recapitulates the numbers of reads for each of the 113 samples and UO per group of samples, respectively. Most rarefaction curves plateaued (except for the three *Culex quinquefasciatus* whole individuals and the two *Aedes aegypti* ovary samples that showed much lower number of reads, as expected) indicating that the sequencing depth was sufficient to describe most of bacterial taxa (Supplementary Note 2 and Supplementary Figure 4).

Whole mosquito bacterial community richness appeared significantly higher for *Aedes aegypti* samples compared to the *Culex* specimens (Figure 1, see Supplementary Note 3 and Supplementary Table 5 for statistical data). In addition, bacterial richness appeared greater for *Culex quinquefasciatus* Slab TC *(Wolbachia*-) compared to the *Culex quinquefasciatus* from the field (*Wolbachia*+) samples. Similarly, bacterial community richness was slightly higher for *Culex pipiens* specimens from the laboratory (Lavar) than from field locations using all three Observed, Chao1 and Shannon diversity indexes. In the case of *Culex* specimen, all samples were positive for *Wolbachia* (Supplementary Note 4), with distinct endosymbiont relative abundance, suggesting a higher bacterial diversity for laboratory specimens as compared to the field ones in this study independent of any previous antibiotic treatment. Noteworthy, *Culex pipiens* samples from the laboratory (Lavar) showed a lower *Wolbachia* relative abundance (Supplementary Figure 5A) yet a higher *Wolbachia* density as compared to field specimen from Camping Europe (Supplementary Figure 5B). These data show differences for *Wolbachia* relative abundance and densities, as confirmed by our statistical analyses (Supplementary Note 4, Supplementary Figure 6).

**Figure 1:**
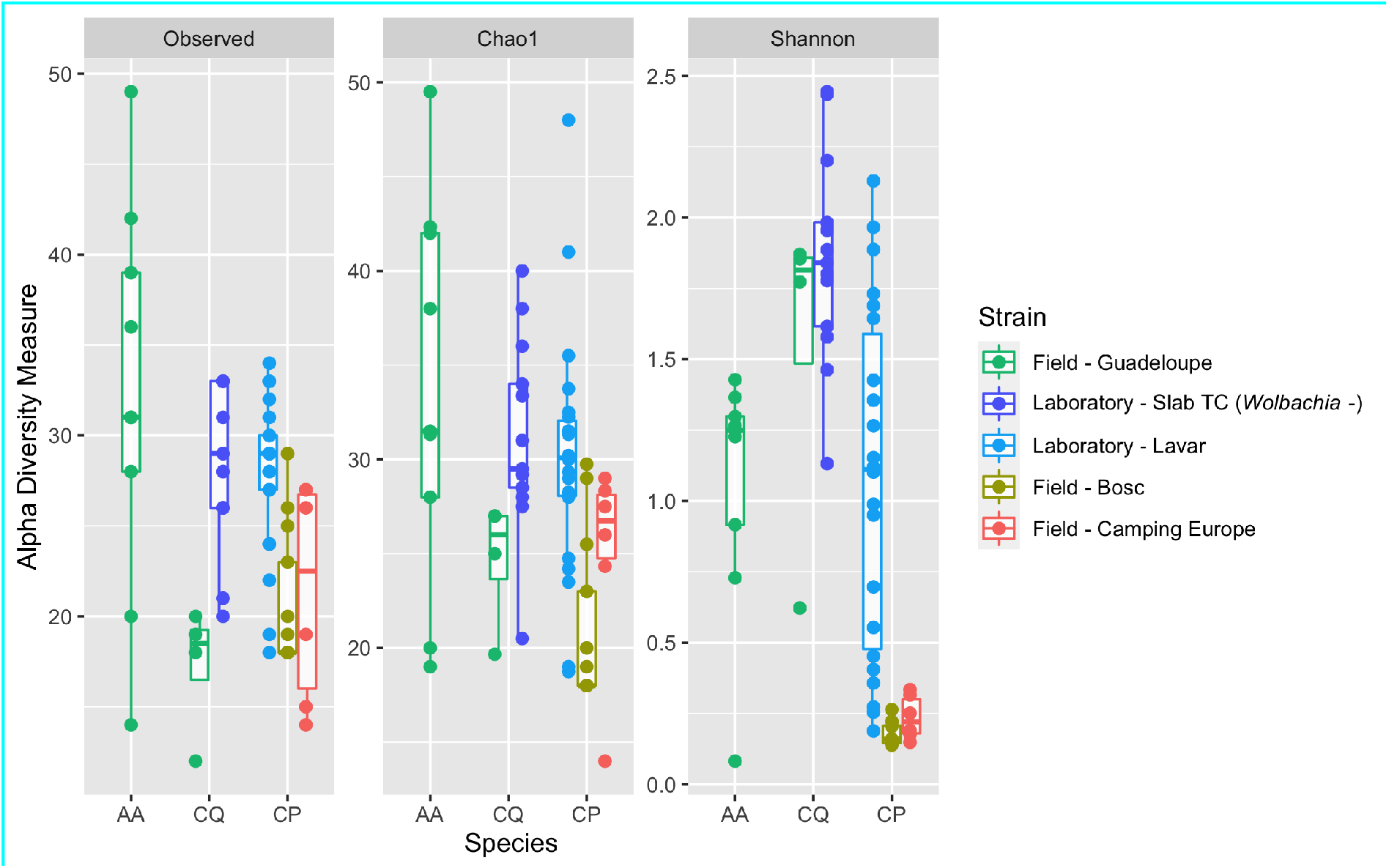
Alpha diversity measures using the Observed, Chao1 and Shannon diversity indexes in whole *Culex pipiens* (CP), *Culex quinquefasciatus* (CQ) and *Aedes aegypti* (AA).

### Species, geography, insectaries conditions as well as *Wolbachia* relative abundance influence mosquito bacterial communities

As expected, *Culex pipiens, Culex quinquefasciatus* and *Aedes aegypti* showed a difference in bacterial community’s composition (Figure 2, Supplementary Figure 7) using NMDS plots, confirming the influence of species on the microbiota. This was statistically confirmed by PERMANOVA (ADONIS, *p*=1e-04, Supplementary Table 2 – Sheet 3). We also observed strong bacterial community differences between *Culex quinquefasciatus* collected from the field in Guadeloupe (*Wolbachia*+) and Slab TC (*Wolbachia*-) (Figure 2, Supplementary Figure 7). This result was confirmed by PERMANOVA that showed a significant p-value (ADONIS, *p*=6e-04, Supplementary Table 2 - Sheet 3) with antibiotic treatment (and consequently null to low *Wolbachia* titers) explaining about 46% of observed variations (ADONIS, R2 = 0.46, see Supplementary Table 2 - Sheet 3). We nevertheless noted the number of *Culex quinquefasciatus* from Guadeloupe (*Wolbachia* +) samples was limited to only 4 samples which comprised a reduced number of reads as compared to the others (<1000 reads), implying a small statistical power for these latter data (Figure 2, Supplementary Figure 7).

**Figure 2:**
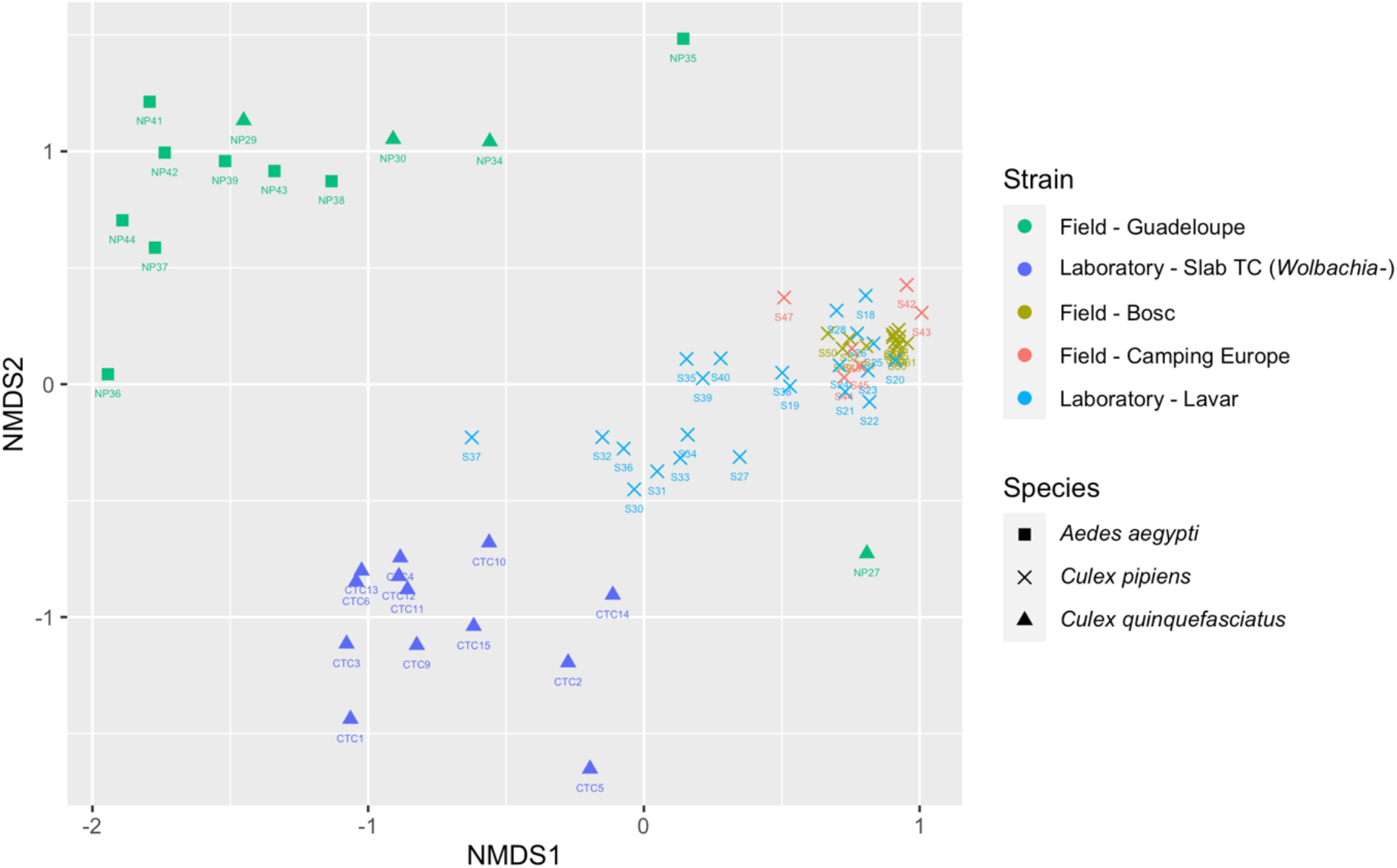
Bray-Curtis based non-metric multidimensional scaling (NMDS) plot of bacterial communities in whole mosquitoes depending on mosquito species and strains

Within *Culex pipiens* specimens, we observed a distinct bacterial community structure in wild samples (Bosc and Camping Europe) compared to the laboratory samples (Lavar), confirming the influence of laboratory setting on bacterial assemblages (Figure 2, Supplementary Figure 7). This difference was partially confirmed by PERMANOVA with a significant difference between bacterial communities from Bosc and Lavar samples (ADONIS, *p*=1e-04, Supplementary Table 2 – Sheet 3), but not between Camping Europe and Lavar samples (ADONIS, *p*=8.3e-02, Supplementary Table 2 – Sheet 3). In addition, we observed a significant difference in bacterial communities of *Culex pipiens* samples from Bosc and Camping Europe (ADONIS, *p*=6e-04, Supplementary Table 2 – Sheet 3), demonstrating the influence of the sampling site on bacterial community structure. *Wolbachia* relative abundance also seemed to influence bacterial community structure in *Culex pipiens* mosquitoes. In fact, we observed a significant difference between microbiota of the *Culex pipiens* mosquitoes (n=41) depending on the *Wolbachia* infection groups (ADONIS, *p*=1e-04, Supplementary Table 2 - Sheet 3) but this effect was also correlated to location (ADONIS, interaction *p*=2e-02, Supplementary Table 2 - Sheet 3).

For the 18 *Culex pipiens* samples for which qPCR has been performed, we confirmed a significant effect of the *Wolbachia* relative abundance on bacterial communities (ADONIS, *p*=1.3e-02, Supplementary Table 2 – Sheet 3) without interaction with any other variable. Nevertheless, we did not observe an influence of the *Wolbachia* density (assessed by qPCR) on mosquito bacterial community structure (Supplementary Table 2 – Sheet 3). These data again reflect differences for *Wolbachia* relative abundance and densities influence, as for alpha diversity measures.

### Mosquitoes harbor multiple symbiotic strains

We identified amplicon sequence variants in our data using unsupervised oligotyping by running it automatically and in some cases manually refining the results for maximum quality assurance (see Materials and Methods). Overall, the 67 unsupervised oligotypes generated by MED were affiliated to a small number of bacteria belonging to 12 different genera: *Wolbachia, Asaia, Legionella, Elizabethkingia, Chryseobacterium, Erwinia, Morganella, Pseudomonas, Delftia, Methylobacterium-Methylorubrum, Serratia* and *Coetzeea*. One unsupervised oligotype (N0318) was only assigned at the order level (Lactobacillales) and represented 0.13% of the total number of reads. A representation of the number of reads per genus is available in Supplementary Figure 8 (panel A).

Whole *Aedes aegypti* individuals, which are known as *Wolbachia* free mosquitoes, individuals were dominated by *Asaia* (85.8%), *Pseudomonas* (10%) and *Chryseobacterium* (3.4%) (Supplementary Figure 8B) while *Culex* spp were dominated by the genus *Wolbachia*. The three most abundant genera in wild *Culex quinquefasciatus* were *Wolbachia* (74.7%), *Asaia* (21.1%) and *Pseudomonas* (2.3%). In *Culex pipiens, Wolbachia* represented 99.8%, 99% and 59.6% of total microbiota in Bosc, Camping Europe and Lavar, respectively. *Culex pipiens* Lavar specimens showed a higher bacterial diversity with *Asaia* as the second most abundant genus (12.7%) and *Elizabethkingia* as the third one (10.1%). On the contrary, *Culex quinquefasciatus* Slab TC (*Wolbachia*-) specimens were dominated by *Elizabethkingia* (35.9%), *Legionella* (29.4%) and *Erwinia* (15%). Surprisingly we observed *Wolbachia* sequences in the latter samples, representing 0 to 10.74% (ie., in sample CTC14, Supplementary Table 3), suggesting potential *Wolbachia* DNA remnants that are not detected through qPCR. As for the reproductive organs, we observed *Wolbachia* dominated both *Culex pipiens* and *Culex quinquefasciatus* samples with 100% relative abundance.

Further, when considering the different genera, we observed several UO affiliated to *Asaia* (N0939, N1147, N1156), *Elizabethkingia* (N0990, N1160), *Chryseobacterium* (N1034, N1035), *Legionella* (N0635, N1065) and *Erwinia* (N0311, N0798) distributed among most mosquito samples (Figure 3). Moreover, while unsupervised oligotype N0711 was the most abundant *Wolbachia* UO, the genus also showed additional variants that were not part of the 15 most abundant UO and were therefore grouped into “Others” (see heatmap with all oligotypes assigned to *Wolbachia* in Supplementary Figure 9). We observed the same *Wolbachia* UO N0711 dominating the ovaries samples, yet co-occurring with additional potential distinct *Wolbachia* variants in these organs too (Supplementary Figure 10).

**Figure 3:**
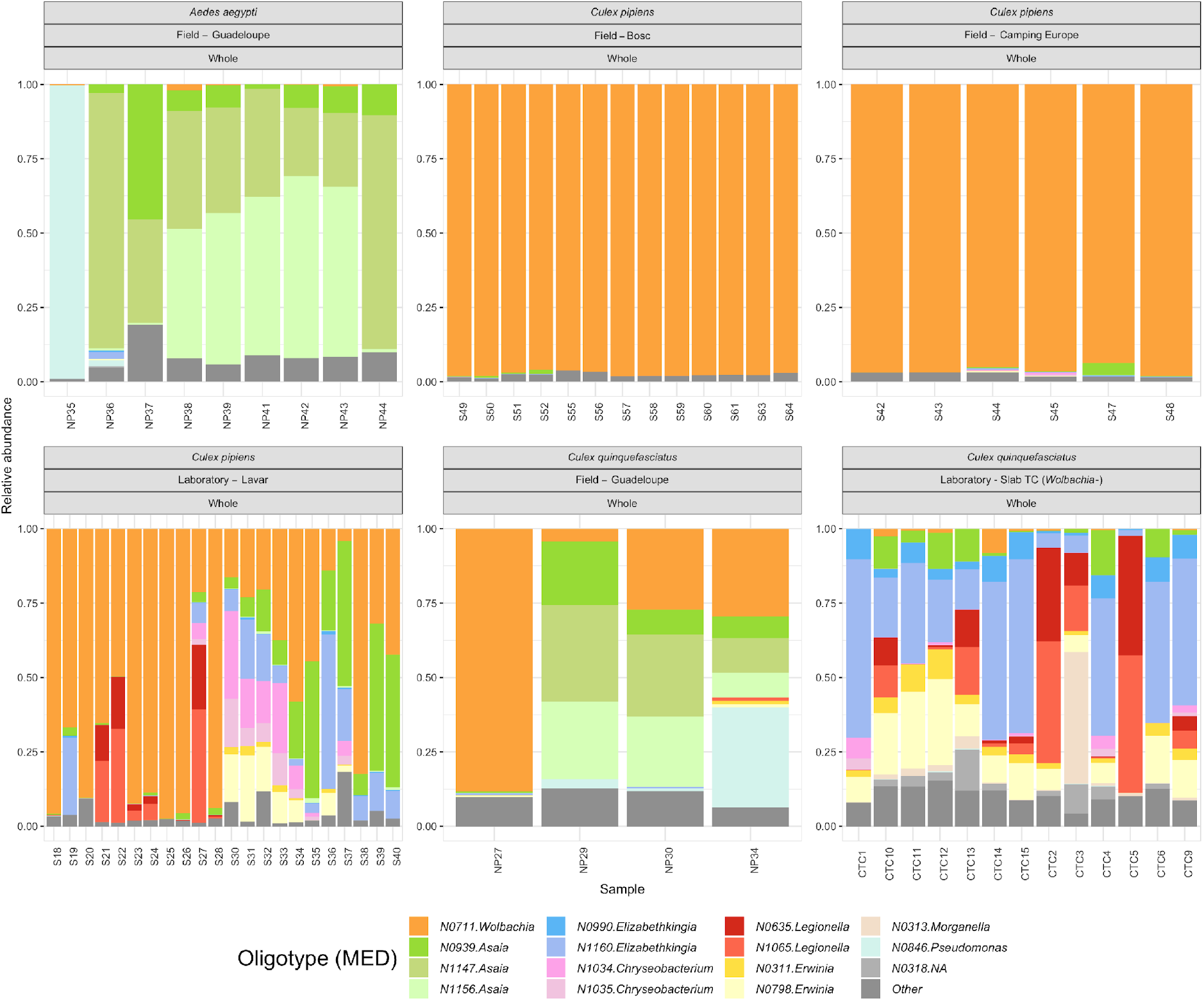
The 15 most abundant unsupervised oligotypes in whole *Culex* spp and *Aedes* individuals in function of mosquito strain. Each color represents an UO with his taxonomic assignment at the genus level. “Other” groups indicate UO that are not included in the 15 most abundant ones. A representation of all UO using a heatmap is available in Supplementary Figure 11.

In order to further investigate the occurrence of putative distinct variants within these mosquito symbionts and commensal taxa, we ran the supervised oligotyping pipeline on genus-level unsupervised oligotypes from genera that showed multiple UO, for both whole individuals and ovary samples.

### Co-occurring and habitat-specific supervised oligotypes

#### *Asaia* supervised oligotypes

*Asaia* supervised oligotypes (SO) were identified within the 1,755,211 sequences affiliated to this genus. We were able to manually decompose it further into 5 supervised oligotypes using 5 nucleotide positions including three abundant and two rare oligotypes. Oligotypes revealed by the entropy found in the entire dataset showed an abundance ratio of ca. 1:2 between the 3^rd^ (376,839 reads) and the two most abundant *Asaia* supervised oligotypes (699,320 and 661,010 reads, respectively). This 1:2:2 ratio between *Asaia* oligotypes suggests that this variation could be explained by a single strain of Asaia with 5 copies of the rRNA operon with subtle differences, consistent with Ribosomal RNA Database^48^ query results for this genus (Supplementary Figure 12, panel C). Yet, breaking down the distribution of SO across individual samples does not support this hypothesis, since the copy number should be preserved across samples. We observed some samples with such expected ratio (e.g., for whole *Aedes* and *Culex quinquefasciatus* mosquito individuals collected in Guadeloupe, a typical sample is NP38) while some other showed distinct ratios like a 5:0:1 (e.g., S50, *Culex pipiens*, Bosc, France), 3:0:1 (e.g., S44, *Culex pipiens*, Camping Europe, France) or 1:0:0 (e.g., S40, *Culex pipiens* isofemelle lines from Lavar laboratory or CTC1, Slab TC, Supplementary Figure 12 panel A1 and A2). These data instead suggest some distinct *Asaia* populations, with one sequence variant found exclusively in Guadeloupe. Interestingly, we observed the co-occurrence of two to three *Asaia* supervised oligotypes in several individual samples. In addition, specimens from insectaries (both Lavar and Slab TC) showed similar *Asaia* SO.

#### *Elizabethkingia* supervised oligotypes

Similarly, we further investigated the *Elizabethkingia* supervised oligotypes by examining the distribution of entropy in 135,187 reads it accounted for. We manually decomposed it further using 1 nucleotide position into two supervised oligotypes; a dominant supervised oligotype C with 124,095 reads and a less abundant supervised oligotype G with 11,092 reads (Supplementary Figure 13). The co-occurrence of both *Elizabethkingia* supervised oligotypes in each individual sample also suggests the presence of closely related *Elizabethkingia* variants. In addition, their differential co-occurrence in almost all samples suggest *Elizabethkingia* variants are not specialized to one or several specific groups of mosquitoes.

#### *Erwinia* supervised oligotypes

Oligotyping 51,363 sequences affiliated to *Erwinia* resulted in 3 supervised oligotypes defined by 8 nucleotide positions. Data showed the presence of one *Erwinia* supervised oligotype (AAGACTTA; 38,025 reads) dominating all samples, as well as two less abundant *Erwinia* variants (TGAGTCGA;7,110 reads AAGACTTG; 4,594 reads, respectively, Supplementary Figure 14). Of note, the Slab TC samples, not dominated by *Wolbachia*, highlighted the co-occurrence of the putative three *Erwinia* variants consistently in all samples, and suggest they could possibly occur in all mosquito samples although they are less or not accessible. These three SO differ from each other by one to 8 nucleotides, representing a minimum of 98,4% percent similarity.

#### *Chryseobacterium* supervised oligotypes

We further investigated the supervised oligotypes of *Chryseobacterium* by studying the distribution of entropy in 106,099 sequences affiliated to this genus. We decomposed *Chryseobacterium* using 3 nucleotide positions into 3 supervised oligotypes. The most abundant *Chryseobacterium* oligotype CGC (66,891 reads) was found only in mosquito samples from Guadeloupe (*Culex quinquefasciatus* and *Aedes*) while oligotypes TAC (12,689 reads) and oligotype TAT (26,519) were observed in samples from France (from the field and lab mosquitoes) as well as in Slab TC (Supplementary Figure 15). Of note, a ratio of ca. 1:2 between the second and the third most abundant SO (12,689 for TAC; 26,519 for TAT) highlighted by the entropy analysis of the whole dataset could suggest they represent two copies of the ribosomal RNA operon of a *Chryseobacterium* bacterium. Such ratio was however not retrieved consistently in all samples, probably discarding this hypothesis. Overall, these data suggest specific *Chryseobacterium* variants in mosquitoes from overseas and continental territories in Guadeloupe and Southern France.

#### *Serratia* and *Legionella* supervised oligotypes

Following the supervised analysis of *Serratia* (8,511 reads), an abundance ratio of ca. 1:2 between the 1st (A; 6,013 reads) and the second most abundant oligotypes G (2498) in almost all samples could suggest that mosquito-associated *Serratia* genomes harbored 2 copies of the rRNA operon (Supplementary Figure 16). Conversely, a ratio of ca. 1:2 between the two most abundant *Legionella* supervised oligotypes CGGA (56,507 reads) and AAAA (34,561 reads) out of 100,773 *Legionella* reads as shown by the entropy found in the entire dataset was not confirmed in all individual samples, suggesting distinct *Legionella* variants (Supplementary Figure 17). Overall, the presence and abundance of *Serratia* and *Legionella* supervised oligotypes varied between samples with no obvious consistent pattern.

#### *Wolbachia* supervised oligotypes

We then further studied the supervised oligotypes *Wolbachia* (4,081,342 sequences), leading to a decomposition into seven oligotypes mainly based on two high Shannon entropy positions 260 and 268 (Supplementary Figure 18, panel C). Therefore, each SO differed by as little as one to two nucleotides (Figure 4). Of note, supervised oligotype CC presented a unique entropy profile with important entropy value for more than 250 positions (data not shown). This particularity can be explained by a deletion at position 192 as we can see in the sequence alignment on Figure 4. *Wolbachia* SO showed a different size (number of reads), with AT accounting for the higher number of reads (3.890.567) and AC accounting the smallest one (2.504, Figure 45). In relative abundance, data showed a dominant *Wolbachia* variants (supervised oligotype AT corresponding to the unsupervised oligotype N0711), accounting for 47 to 100% and rare “variants” that represented between 0 to 26% for supervised oligotype AG, 0 to 21% for supervised oligotype GT, 0 to 16% for supervised oligotype TT, 0 to 0.5% for supervised oligotype CC and 0 to 0.25% in the different studied samples (Supplementary Table 6 - Sheet 1).

**Figure 4:**
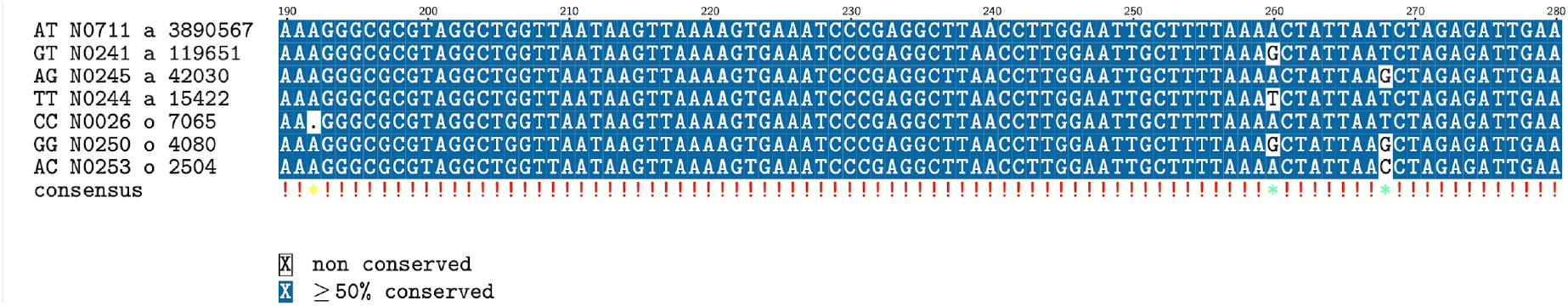
Alignment of the seven *Wolbachia* supervised oligotypes retrieved in this study. The first letters correspond to the supervised oligotype (at the 260 and 268 positions that varied), the first number represents the corresponding unsupervised oligotype, the following letter indicates abundance information (“a” means that unsupervised oligotype is among the 15 most abundant, “o” means that unsupervised oligotype is among “Other” in Figure 3), and the last number indicates the size of supervised oligotype (number of reads).

When studying the distribution of these supervised oligotypes, we observed the co-occurrence of one to four different SO in a single sample like in S42 or CTC6, respectively (Supplementary Figure 18, panel A1 and A2). We also noted multiple co-existing SO in samples with low (Slab TC) to medium or high (most of *Culex pipiens* from Bosc locality for instance) *Wolbachia* relative abundance, suggesting the presence of these supervised oligotypes is not linked to the *Wolbachia* predominance.

We eventually blasted these supervised oligotypes in order to investigate the presence of these putative variants in the available databases. On the one hand, we observed the dominant supervised oligotype AT had a match with 100% identity with *Wolbachia pipientis* isolates including TuFIVIA19m (NCBI Accession number MN123175.1) while GT, AG, CC, TT, and GG had a match with 99.5% to 99.75% identity to the same sequence. On the other hand, supervised oligotype AC had a match with 100% identity to three distinct *Wolbachia* strains including *Wolbachia* endosymbiont of *Acrocephalus palustris* isolate Acr172 (NCBI Accession number MF374624.1), *Wolbachia* endosymbiont of *Ectropis obliqua* clone WX-03 (NCBI Accession number KU058642.1), *Wolbachia* endosymbiont of *Medythia nigrobilineata* strain Mni (NCBI Accession number GU236941.1). While the sequence variants that occur in a small number of positions throughout the amplicons give credence to true biological diversity, it is difficult to exclude PCR amplification-associated systematic errors to result in abundant variants, especially in the presence of a remarkably numerous template.

### Metagenome screening of *Wolbachia* supervised oligotypes

Since amplification-free shotgun metagenomes are less prone to PCR errors, we further investigated the presence of additional *Wolbachia* variants suggested by the highly-resolved analysis of amplicon sequences in single individuals using metagenomic datasets generated independently. Using variable regions of supervised oligotype sequences as ‘primers’ we searched high-quality metagenomic short reads in available *Culex pipiens* ovary metagenomes^30^ and egg-rafts metagenomes^47^ from France and different locations in Northern Africa generated using the Illumina sequencing technology. For that purpose, we extracted small nucleotide fragments (Supplementary Figure 19) including the two highest entropy positions and searched for their presence in the *Culex* metagenome datasets. As expected, supervised oligotypes AT counted a huge number of hits within the four *Culex pipiens* ovary metagenome samples, that is between 96 to 333 hits (Supplementary Figure 20). In addition, other supervised oligotypes showed single hits (Supplementary Figure 20) which could further suggest putative multiple *Wolbachia* infection in *Culex* samples, although these remain based on very small read numbers and therefore do represent ambiguous data.

## DISCUSSION

In this study, we realized a fine-scale 16S rRNA gene analysis of the bacterial communities of *Culex pipiens, Culex quinquefasciatus* and *Aedes aegypti* mosquito individuals collected in continental and overseas regions of France. We examined the bacterial assemblages in the whole bodies as well as dissected ovaries from specimen collected in the wild or in insectaries (Lavar *C*.*pipiens* and antibiotically-treated *C. quinquefasciatus* Slab TC) as well as with low to high *Wolbachia* relative abundance and densities. We used Minimum Entropy Decomposition (MED) and oligotyping that use the Shannon entropy to identify ecologically relevant and to discriminate taxa that differ by as little as a single nucleotide in the sequenced region of these datasets^24,25^.

### The influence of host species, geography, environmental conditions, symbiont relative and absolute abundance on mosquito microbiota

Bacterial community structure differed significantly between our *Culex pipiens, Culex quinquefasciatus* and *Aedes aegypti* samples, confirming different mosquito microbiota for each species. Although bacterial communities of wild *Aedes* specimens were dominated by *Asaia* taxa (85.8%), *C. pipiens* and *C. quinquefasciatus* individuals were dominated by *Wolbachia, Asaia* and *Elizabethkingia*, yet in different proportions for each species. The predominance of *Wolbachia* was in agreement with previous studies in *Culex pipiens*^49–51^ and *Culex quinquefasciatus*^52,53^. Similarly, *Asaia* is one of the most-frequently detected genera in *Aedes aegypti* microbiota^54–58^, although it was found in lower abundances than in our study. It is likely that the predominance of *Wolbachia, Asaia* and *Elizabethkingia* in specimens from both *Culex* species explains the observed partial overlap in the overall community structure of both species. Geography at a fine scale (such as Bosc and Camping l’Europe sites which are not far from each other) was nevertheless shown to influence bacterial communities within *Culex pipiens*. These data were in agreement with previous studies, showing the site of mosquito collection influences bacterial community structure at a large geographic scale^59^.

We then observed clear differences between field and insectaries mosquito microbiota. Drastically distinct bacterial profiles in individuals from *C. quinquefasciatus* lines which were treated (at least five generations before sampling) and not treated with antibiotics was not a surprise, as antibiotics are commonly used to eliminate resident microbiota from mosquitoes. Nevertheless, *C. pipiens* specimens reared in the laboratory for several consecutive years (Lavar strain) unexpectedly showed a higher bacterial richness as compared to wild mosquito individuals. These data differed from previous studies, showing a particularly poor bacterial richness and diversity in laboratory mosquitoes in *Anopheles* species^60^. One hypothesis could be that selection on *Culex* microbiota composition could be stronger in environmental conditions while it might be more relaxed in the lab, where mosquitoes are fed every day and competition is lower allowing the presence of diverse bacteria. Another hypothesis could be that slightly lower *Wolbachia* relative abundance in *Culex* specimen from the lab, as observed herein, could allow for a greater bacterial diversity to be observed (Supplementary Figure 5 A). Nevertheless, *Wolbachia* densities as detected by qPCR were noteworthy higher in *Culex pipiens* from the lab (Supplementary Figure 5B).

These data underlines high-throughput sequencing indicates relative abundances of bacterial taxa, and cannot be directly linked to absolute abundance, in agreement with Williamson and colleagues^61^. In addition, it suggests high bacterial diversity in the lab could possibly be associated with high bacterial loads for the remaining taxon. These different hypotheses might not be exclusive from one another and future studies should investigate these scenarios using more samples and laboratory conditions. In any case, bacterial community differences highlight the importance of lab vs. environmental conditions on microbiota and reminds us to remain cautious and critical when conducting experimental infection using laboratory strains. Mosquito microbiota from lab strains could indeed respond differently to pathogens in experimental conditions and not mimic wild mosquito individuals.

### Closely related 16S rRNA gene sequences highlight multiple symbiont variants

Overall, our combined unsupervised and supervised oligotyping analyses revealed that some genus-level mosquito-associated bacterial taxa contain putative previously undetected co-occurring variants at the scale of single individuals. In particular, it highlighted the predominance of certain variants yet the presence of additional but less abundant ones that would not have been seen using a classical clustering approach.

We noted the co-existence of closely related *Elizabethkingia* and *Erwinia* variants in several individuals, suggesting the possibility of some gene content variation and functional differences, warranting further analyses at the genome scale. The bacterium *Elizabethkingia anophelis* was first isolated from *Anopheles* mosquitoes in 2011^62^ and later detected in *Aedes* dengue fever vector mosquitoes but not in *Culex*^63^. *Elizabethkingia*-like bacteria were identified as interesting candidate for paratransgenic approaches due to some strong anti-*Plasmodium* activity. However, a recent study showed a single gene disruption in one *Elizabethkingia anophelis* strain was at the origin of an atypical mutation rate and a severe human disease outbreak in the US from 2015 to 2016^64^. Although a 1% nucleotide distance separated the outbreak strain from its closest environmental *E. anophelis* relatives at the whole genome level, the authors^64^ showed the integration of a mobile genetic element in a specific gene within this species can have serious implications. In line with this, the co-existence of closely related *Erwinia* variants showing more 98,4% percent similarity raise questions about the potential phenotypic differences of each strain. Members of the *Erwinia* group are phytopathogenic and exploit insects both as hosts and vectors^65^. The presence of a dual *Erwinia* infection might have implication for crop managements, suggesting some monitoring and further genome-scale analyses are needed.

In addition to different strains coexisting in a single individual, our study showed some variants specific to certain populations of mosquitoes, like for *Asaia* and *Chryseobacterium* in Guadeloupe vs. continental France. We actually observed one *Asaia* variant that seemed to occur in Guadeloupe only, and not in French populations. These data are in agreement with previous studies, showing that different *Asaia* strains are present in different mosquito populations^66^. The authors indicate the importance of uncovering the diversity of *Asaia* strains, proposed as potential antiplasmodial agents, in the context of paratransgenic approaches. Similarly, the observation of specific *Chryseobacterium* sp. strains in different geographic locations is in agreement with their mutualist/ commensal characteristics. *Chryseobacterium culicis sp. nov*., was first isolated from the midgut of the mosquito *Culex quinquefasciatus*^67^ and later on described as having a beneficial role for their host, rescuing axenic larval mosquito development^17,63^, while inducing a weak immune response. Previous work showed this bacterial species is well adapted to life in the gut of insect^68^ and one can expect high level of specificity from these mutualistic relationships.

Further, our data suggest the putative co-occurrence of several *Wolbachia* oligotypes in a single *Culex* specimen. Although systematic PCR errors in samples dominated by one or few bacterial taxa could influence the observed results, we retrieved the same oligotyping pattern for wild *Culex pipiens* individuals that have high to low *Wolbachia* densities, and that all together offer a *Wolbachia* relative abundance gradient. On the one hand, these data could putatively suggest possible co-infection patterns in independent samples and with no link to *Wolbachia* densities. On the other hand, the presence of varying *Wolbachia* 16s rRNA gene sequences within a single mosquito individual could not be confirmed with more than one hit when screening seven different *C. pipiens* ovary and egg-rafts metagenomes collected in Southern France and Northern Africa (Supplementary Figure 20) avoiding any conclusive remark at this point. Nevertheless, the very low number of AC sequences retrieved in the metabarcoding datasets (ca. 2,500) suggests they might be too rare to be detected in the available Illumina metagenomes datasets. In addition, the detection of some *Wolbachia* reads in few *Culex* Slab TC sample was not expected and might be due to some DNA remnants rather than true bacterial cells or Horizontal Gene Transfer (HGT) events. Although HGT of functional genes between *Wolbachia* and its host has been reported^69^, no 16S rRNA genes HGT has been reported in *Wolbachia* to our knowledge. More generally, 16S rRNA genes HGT seems a rare event occurring at the intra-genus and intra-species levels^70^.

The 100% Blast matches with *Wolbachia* sequences for both AT and AC supervised oligotypes add more weight on the likelihood of having an “A” at the first high-variation position (260), while the second position (268) is possibly more variable. In other words, it reinforces the probability of a biological variation at position 268 while it diminishes the one at position 260. In line with this observation, we noted a homopolymer region before the first high entropy position “A”, which could favor a possible sequencing or polymerase error while the second high-variation position variation remains independent of this region and might be biologically true. Although this hypothesis could not be verified here, these data would suggest there are at least two *Wolbachia* variants (represented by supervised oligotypes AT and AC) while the remaining supervised oligotypes GT, AG and CC possibly represent artefactual variations due to sequencing errors or polymerase errors. These two potential *Wolbachia* variants differed by one nucleotide, representing only 0.2% variation for a 428 bp fragment. These subtle nucleotide variations might have been obscured using OTU clustering at 97% or 99% in high-throughput sequencing and analysis, and therefore remained undetected till now. The presence of one nucleotide position variation at the 16SrRNA gene level could nevertheless suggest some larger differences at the genome level. These data are congruent with our previous analysis showing 168 to 612 Single Nucleotide Variants (SNV) for each of the four *Wolbachia* Metagenome-Assembled Genomes (MAGs) reconstructed from the ovaries metagenomes^30^. Of note, these values might be underestimated since the different MAGs were not complete, showing between 93 and 127 contigs. In particular, MAGs were missing some of the highly-variable phage sequences that could not be assembled due to repetitive sequences leading to assembly breaks. These data could also indicate a horizontal transmission of rare *Wolbachia* variants.

These results nevertheless differ from previous studies showing a unique *Wolbachia* 16S rRNA sequence and the lack of single nucleotide polymorphism in distinct *C. pipiens* populations from different geographic localities distances using Sanger sequencing^71^. However, other studies have been showing high levels of multiple *Wolbachia* infection in other insects like the ant *Formica exsect*a^72^, with the hosts harboring up to five distinct *Wolbachia* strains. Authors revealed a maximum of 3% variations in the highly variable *wsp* gene, indicating highly divergent *Wolbachia* strains for this species. In addition, Arai and colleagues^73^ also found a triple *Wolbachia* infection in tea pest *Homona magnanima*. The authors established uninfected and singly infected lines using antibiotics and revealed distinct CI intensities and/or mutualistic effects between the different *Wolbachia* lines^73^. Although the putative *Wolbachia* variants detected here differed by much less nucleotides, it remains unknown whether they may have distinct roles and induce distinct host-phenotype. In particular, a possible divergent impact on the biology of mosquito species including susceptibility to pathogens such as WNV (West Nile virus) for *Culex* species remains unknown. Related to these issues, the presence of dominant and rare *Wolbachia* variants within the same individual would also suggest caution is needed in programs of *Wolbachia* transinfection. Overall, we believe the two supervised oligotypes AT and AC are worth to be noted, although the biological significance of these *Wolbachia* sequences is a question for further study.

### The importance of accounting for both bacterial relative and absolute abundance in fine-scale microbiota analysis

In general, this study confirms that the dominance of *Wolbachia* on most mosquito samples might have hampered estimating the actual diversity of symbionts associated with mosquitoes in previous studies. qPCR analysis targets the relative abundance of *Wolbachia* per host copy number and presumably better represents the actual *Wolbachia* titer. However, it might not be sensitive enough to distinguish and account for closely related and rare variants detected through high-throughput sequencing. As expected, analyzing set of mosquito samples with supposedly no to very low *Wolbachia* abundance (*e*.*g*. Slab TC) and therefore not dominated by a single bacteria, allowed to shed light on clear patterns of closely related variants co-occurring in mosquito specimens, like for *Elizabethkingia* and *Erwinia*. Although the *Elizabethkingia* variants differed by a single nucleotide at the 16S rRNA gene amplicon-level, they indeed might not be functionally redundant. The presence of closely related symbionts in multicellular hosts is now commonly reported, with several studies reporting some generalized niche partitioning between symbionts. Brochet and colleagues^74^ showed niche partitioning allowed the co-existence of closely related bacteria in the gut of honey bees. Similarly, functional diversity was shown to enable the coexistence of multiple strains and niche partitioning in deep-sea *Bathymodiolus* mussels^75,76^ or in the *Rimicaris exoculata* holobiont^77^. Although transovarian transmission of symbionts is more common in arthropod while horizontal transmission is usually observed in marine environments, niche partitioning and functional diversity might allow symbiosis to sustain itself together with its host resilience and evolution.

## Conclusion

Overall, this study allowed a high-resolution characterization of mosquito microbiota and suggests that co-infection pattern of symbionts could be common in mosquito species. Future studies on the maintenance, and importance of bacterial variation at the genomic-scale will allow understanding the possible phenotypic effect of multiple infection in mosquitoes as well as the evolutionary forces leading to this pattern. In addition, the less-abundant and neglected members of the mosquito bacterial microbiota may have possible divergent history traits and functions that require further genome-resolved metagenomic, microdiversity analysis and strain comparison.

## Code availability

A reproducible bioinformatics workflow including, scripts and supplementary materials used for this study is available at https://github.com/jreveillaud/Mosquito-Symbiont-Microdiversity.

## Data availability

Raw sequencing data are available through the European Nucleotide Archive (PRJEB43079).

## Acknowledgments

We thank Frederic Mahé and Renata Servan de Almeida for their insights and help with computational and logistic matters, respectively. We are also thankful to Sandra Unal for insectories maintenance. This work was supported by the ERC RosaLind starting grant to J.R, the INRAe Animal Health Department and Occitany Region to H. S, the EU project MALIN and the Guadeloupe Regional Council under the European Research and Development Funds (ERDF) 2014-2020 program (Grant 2018-FED-1084) to NP.

