## Supplementary Material for "The mosquito microbiome includes habitat-specific but rare symbionts"

**Supplementary Note 1. Quality filtering and decontamination of raw Illumina reads**
We obtained 6,802,607 high-quality sequences after stringent quality filtering of Illumina reads from the 136 mosquito samples, with an average number of 50,019 sequences per sample. We removed samples that counted less than 1,000 reads except 5 samples of *Culex quinquefasciatus* and *Aedes* from Guadeloupe (NP20, NP36, NP29, NP30, NP34) to ensure representation of both species from Guadeloupe (Supplementary Table 1 sheet 1). Discarded samples included 4 *Culex pipiens* specimens, 2 *Aedes aegypti* specimens and 7 blanks (See Supplementary Table 1 for details). The Minimum Entropy Decomposition (MED) method generated 70 representative unsupervised oligotypes (Supplementary Table 1 sheet 2) that counted 6,487,244 reads after discarding 313,778 reads because of default filters application. We then removed three contaminant unsupervised oligotypes (N0005 - *Enhydrobacter*, N0852 – *Pseudomonas* and N0315 - *Rahnella1*) that were more abundant in blanks than in true samples after Decontam analysis (17,460 reads). The identified bacteria could be part of the so called kitome as both *Enhydrobacter* and *Pseudomonas* are common contaminant genera detected in sequenced negative `blank' controls. Because of the removal of these contaminant unsupervised oligotypes, two true samples counted less than 1000 reads and were thereafter removed from the dataset (Supplementary Table 1). We eventually removed the remaining 8 blanks (16,211 reads) for downstream analyses. These series of filters resulted in a total of 113 samples represented by 6,452,623 high-quality reads that clustered into 67 unsupervised oligotypes.

**Supplementary Note 2. Rarefaction curves**. Most rarefaction curves reached a plateau, suggesting a sufficient sequencing depth to describe most of bacterial taxa. Only few of them increased steadily indicating a higher sequencing would be needed, notably for the samples from *Culex quinquefasciatus* (whole individuals) and *Aedes aegypti* (ovary) collected in Guadeloupe (Supplementary Figure 4). We nevertheless retained the latter samples for downstream analyses in order to have a representation of mosquito individuals from Guadeloupe*.* We compared alpha and beta diversity analysis including and excluding them to ensure they had no influence on the remaining analyses (data not shown).

**Supplementary Note 3. Bacterial community richness.**
***Aedes* vs. *Culex* spp**. We observed a higher average value of alpha diversity within wild *Aedes aegypti* (mean of the Observed index=32.2; mean of the Chao1 index=33.5; mean of the Shannon index=1.06, Supplementary Table 5) compared to wild *Culex pipiens* (mean of the Observed index=20.8; mean of the Chao1 index=22.2; mean of the Shannon index=0.2, Supplementary Table 5) and wild *Culex quinquefasciatus* (mean of the Observed index=17.3; mean of the Chao1 index=24.7; mean of the Shannon index=1.53, Supplementary Table 5). This difference was significantly supported by ANOVA (p.val for Observed=3.6E-04; Chao1=1.2E-03; Shannon=9.09E-10) for each diversity index and Tukey's test for two indexes (see Supplementary Table 2 - Sheet 2).

***Culex quinquefasciatus***. Slab TC individuals had a mean of species diversity about 28.4, 31.1 and 1.85 for the Observed, Chao1 and Shannon indexes, respectively, while *Culex quinquefasciatus* from the field (*Wolbachia*+ samples) showed ones of 17.2, 24.7 and 1.53

(see Supplementary Table 5). These differences were partially confirmed by ANOVA (p.val for Observed=3.2E-04; Chao1=3.6E-02; Shannon=0.2 NS; (Supplementary Table 2, sheet 2).

***Culex pipiens.*** We observed within *Culex pipiens* from Lavar an average diversity value of 27.9, 30 and 1.1 for the Observed, Chao1 and Shannon indexes, while *Culex pipiens* from Bosc and Camping Europe had ones of 20.6, 20.9 and 0.18, and 21.3, 24.9 and 0.24, respectively (Supplementary Table 5). These differences were statistically confirmed by ANOVA (p.val for Observed=4.5E-05; Chao1=2.4E-04; Shannon=2E-06) and Tukey's test that showed a significant difference between *Culex pipiens* from Lavar and *Culex pipiens* from Bosc and Camping Europe (Supplementary Table 2 - Sheet 2).

Noteworthy, species diversity estimates obtained with Observed and Chao1 appeared as relatively similar, while a distinct trend was obtained using the Shannon index. We observed a slight bacterial diversity increase for *Culex quinquefasciatus* individuals collected in the field and for Slab TC (*Wolbachia*-) specimens, and a decrease for *Aedes aegypti* using the Shannon index as compared to with Observed and Chao1 indexes. This pattern could be due to the fact that few species are making up a large number of the community in the latter samples (reflecting inequality).

**Supplementary Note 4. *Wolbachia* relative abundance and density**. As expected, the Slab TC (treated by Tetracycline) samples showed 0 to very low DNA quantity of *Wolbachia* (i.e., CTC12) quantified by qPCR and corresponded to the "Low infection" groups identified using HCA with only one outlier (CTC14, "Medium infection" sample, 0.1074139, Supplementary Table 3 – Sheet 2, *Wolbachia*- samples). The remaining samples showed some DNA quantity > 0.1, and were verified as *Wolbachia*+ by qPCR. However, the "Medium infection" and "High infection" from the HCA based on relative abundance of *Wolbachia* only showed a partial correspondence with the qPCR results (Supplementary Table 3 – Sheet 2 and Supplementary Figure 6). These discrepancies were expected and can be explained by the comparison of two distinct metrics: raw abundance of *Wolbachia* (qPCR) and relative abundance of *Wolbachia* (used for HCA). We nevertheless retained the 3 HCA defined groups in order to investigate *Culex* bacterial community structure in function of *Wolbachia* relative abundance in downstream analyses. Of note, only one *Wolbachia*+ sample (NP29, *Culex quinquefasciatus* from Guadeloupe) showed a relatively high value using qPCR although it was assigned to the "Low infection" group (3.48, Supplementary Table 3 – Sheet 2). In addition, *Culex pipiens* specimen from the lab showed a higher *Wolbachia* load measured with qPCR as compared to field ones (Supplementary Figure 5).

**Supplementary Table 1**. Sheet 1. Samples collected with sequenced Sample ID, Mosquito strain, Field vs. lab, Country, Type, species, Collection date, MiSeq Run, True vs. Control sample, Read Depth. Sheet 2. Raw count table of the 123 retained samples after MED analysis and removing samples with less than 1000 reads (except 5 *Culex quinquefasciatus* and *Aedes* samples from Guadeloupe NP20, NP36, NP29, NP30, NP34). Sheet 3. Final count table of the 113 samples retained after Decontam analysis. Sheet 4. Taxonomy table of the 70 unsupervised oligotypes before Decontam analysis. Sheet 5. Final taxonomy table of the 67 unsupervised oligotypes. A * indicates samples that were removed from the analysis dataset of a number of reads inferior to 1000. A + indicates samples removed after Decontam analysis.

**Supplementary Table 2**. Alpha and Beta-diversity statistical analyses. Sheet1: Estimating influence of sequencing depth on alpha-diversity using ANOVA. Sheet 2: Estimating Alpha-diversity within groups using ANOVA and tukey's tests. Sheet 3. Beta-diversity using PERMANOVA.

**Supplementary Table 3**. Sheet 1: Statistics of *Wolbachia* infection within groups obtained with HCA analysis from all samples (n=123). Sheet 2: *Wolbachia* load measured by qPCR in 34 samples (qPCR_Ratio_Target_Ref column) and percent of *Wolbachia* in these samples with their *Wolbachia* infection group computed by HCA.

**Supplementary Table 4**. Number of samples, reads and unsupervised oligotypes by species, location and organ.

**Supplementary Table 5.** Mean of alpha diversity indexes in whole mosquitoes by species and location.

**Supplementary Table 6.** Relative abundance of the supervised oligotypes within samples. A * indicates supervised oligotypes that matched with unsupervised oligotypes post-MED analysis. Sheet 1. *Wolbachia*, Sheet 2. *Asaia*, Sheet 3. *Elizabethkingia*, Sheet 4. *Erwinia*, Sheet 5. *Chryseobacterium*, Sheet 6. *Serratia*, Sheet 7. *Legionella*.

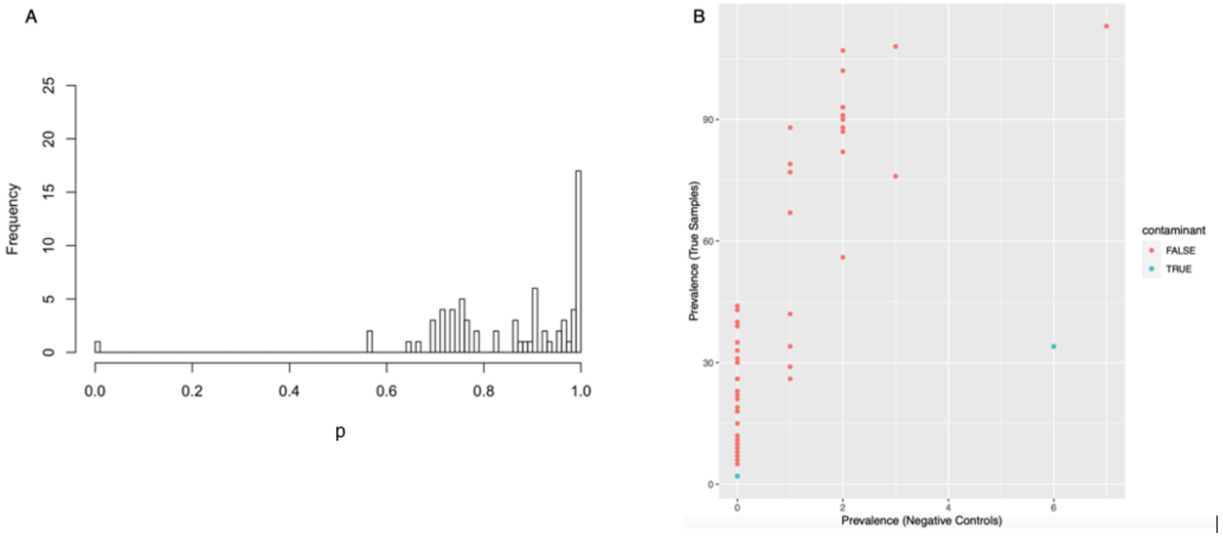


**Supplementary Figure 1**. Decontam results. A. Histogram generated with frequency of unsupervised oligotypes in function of p (probability, also referred to score, used for classifying contaminants in Decontam using the prevalence-based contaminant identification), with contaminant = TRUE if p is identified at 0.65 herein. B. Prevalence of true samples in function of prevalence of negative controls.

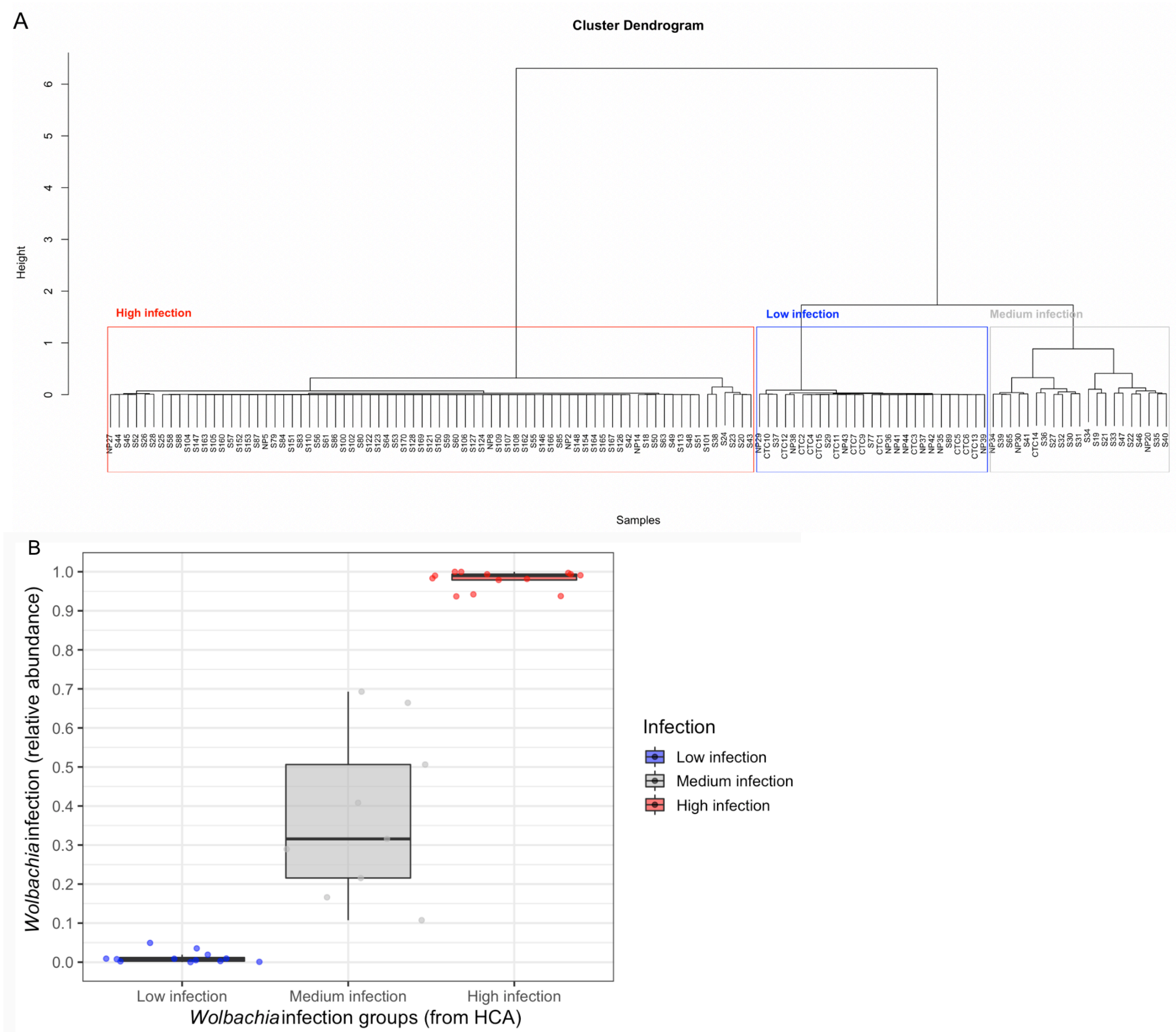


**Supplementary Figure 2:** A. Cluster dendrogram from the HCA analysis based on the distance matrice of *Wolbachia* relative abundance in all samples (n=123). Height reflects the distance between the samples/clusters. B. Boxplot of *Wolbachia* infection (relative abundance) by *Wolbachia* infection groups (from HCA).

.

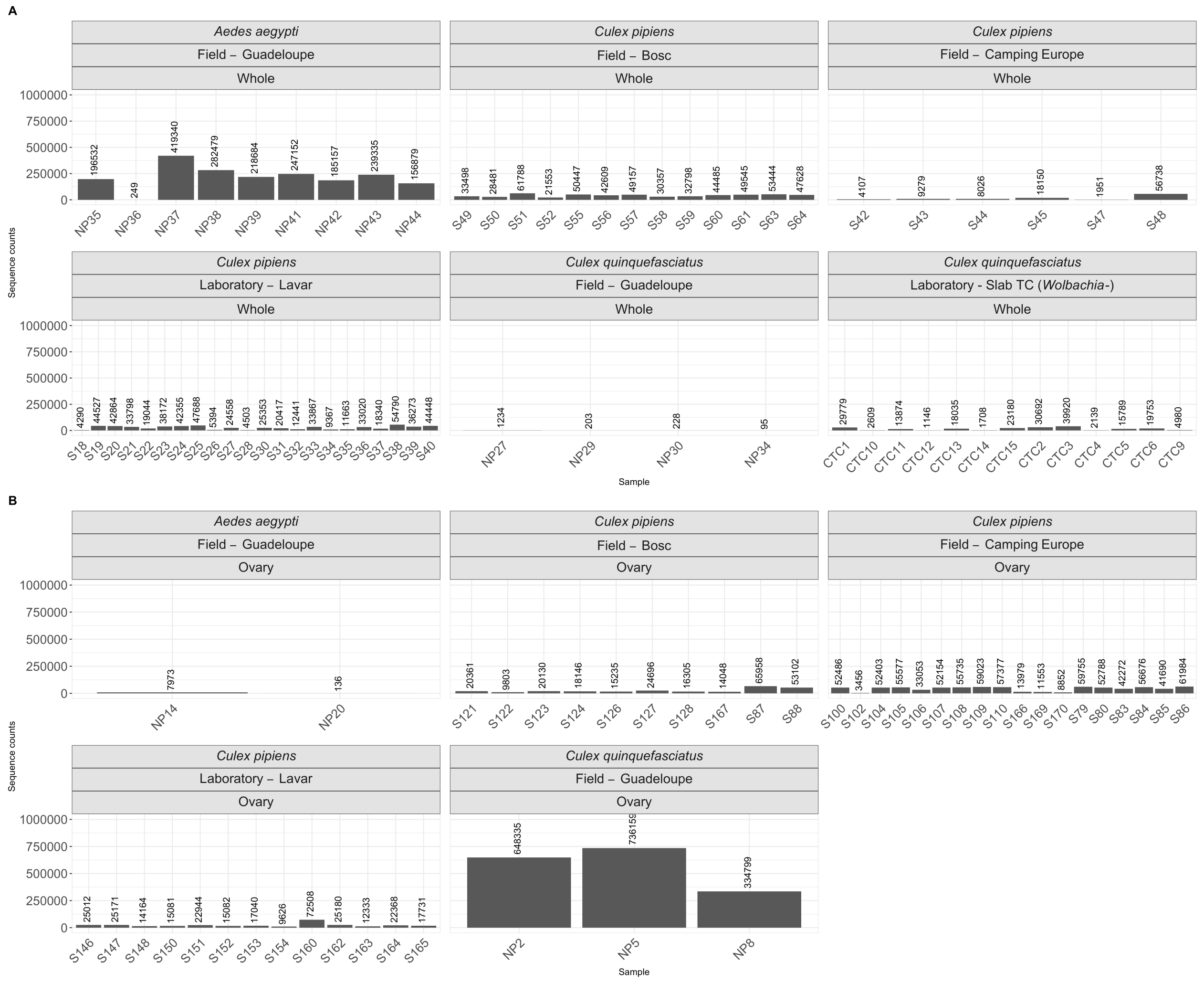

**Supplementary Figure 3**: Number of reads for each of the 113 samples retained after MED analysis, decontam and the removing of samples with less than 1000 reads (with the exception of NP20, NP36, NP29, NP30, NP34 as detailed in Supplementary Note 1). A=Whole and B=Ovary.

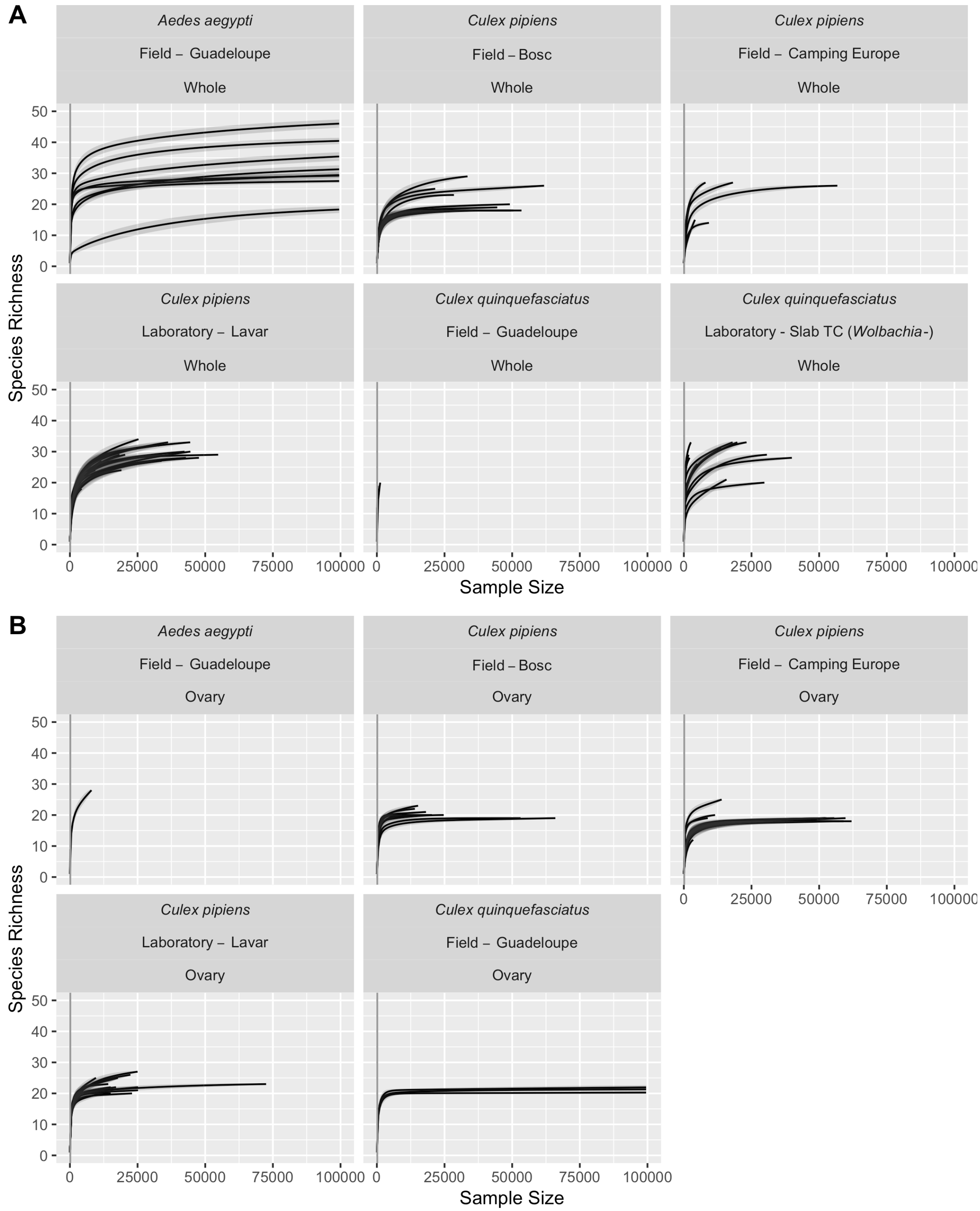


**Supplementary Figure 4**: Rarefaction curves of whole (A) and ovary (B) samples from *Culex pipiens*, *Culex quinquefasciatus* and *Aedes aegypti* samples.

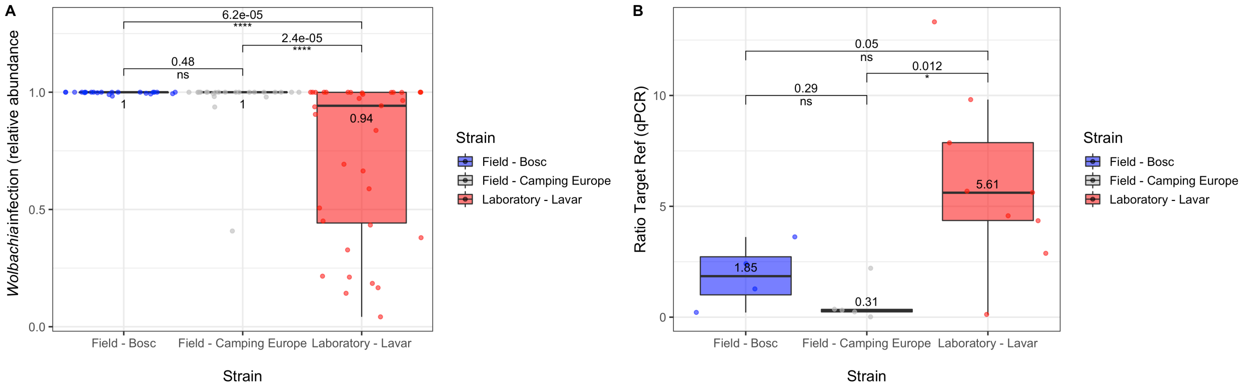


**Supplementary Figure 5:** A: Boxplot of *Wolbachia* infection (relative abundance) in *Culex pipiens* strains. B: Boxplot of *Wolbachia* density (Ratio Target Ref from qPCR) in *Culex pipiens* strains*.* Line in boxplot represents the median. Paired Wilcox-test p-values and significance symbols (ns: p > 0.05, *: p <= 0.05, **: p <= 0.01, ***: p <= 0.001, ****: p <= 0.0001) are reported into brackets.

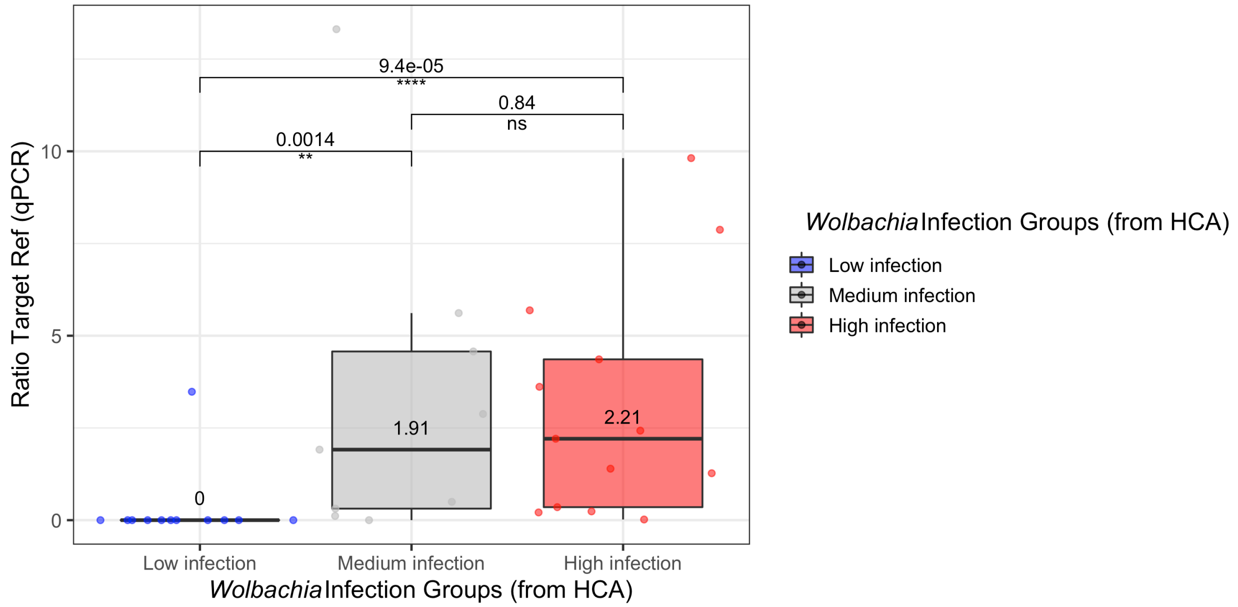


**Supplementary Figure 6:** Boxplot of *Wolbachia* density (qPCR) for each *Wolbachia* infection groups formed by the HCA analysis of *Wolbachia* relative abundance in *Culex* spp. Dots correspond to *Wolbachia* density determined by qPCR. Line in boxplot represents the median of Ratio Target Ref (qPCR) by *Wolbachia* infection group. Paired Wilcox-test p-values and significance symbols (ns: p > 0.05, *: p <= 0.05, **: p <= 0.01, ***: p <= 0.001, ****: p <= 0.0001) are reported into brackets.

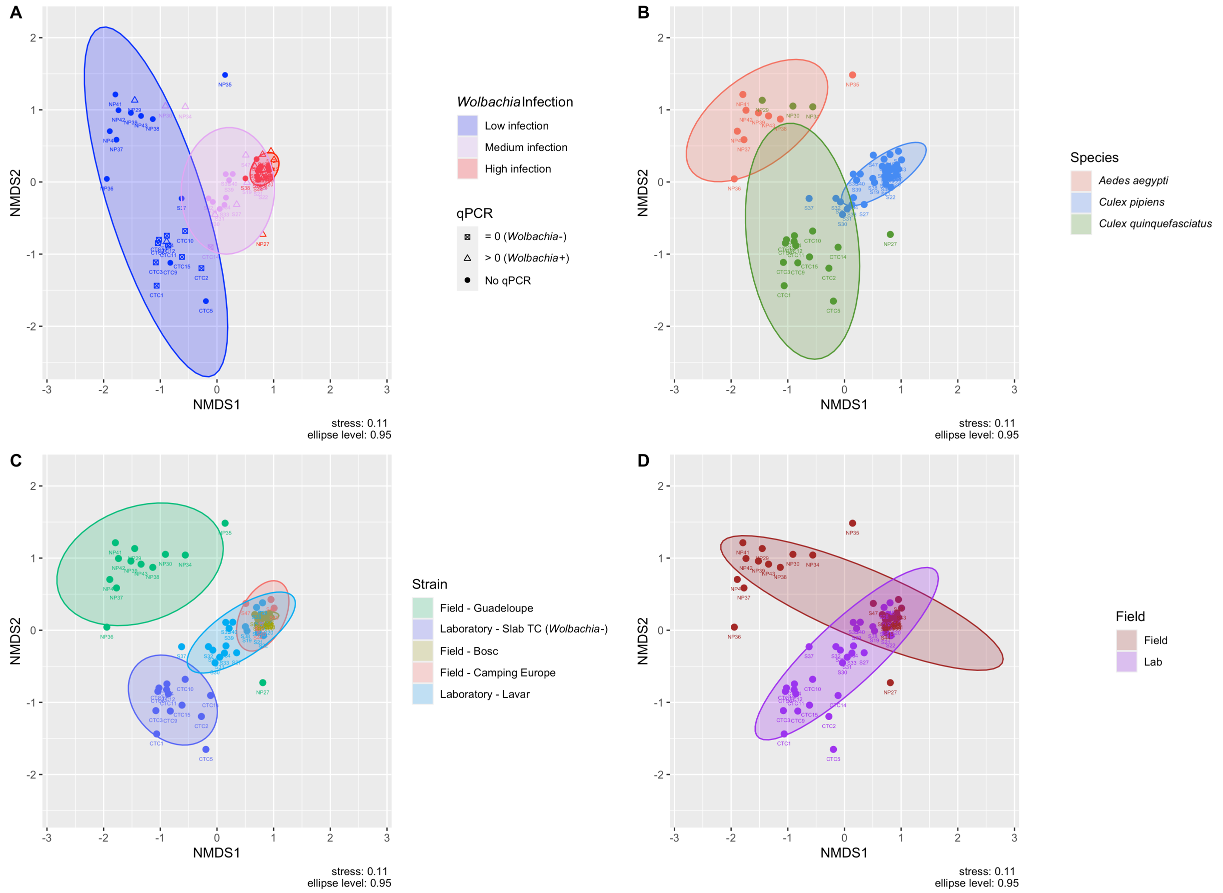


**Supplementary Figure 7**: Bray-Curtis based non-metric multidimensional scaling (NMDS) plot of bacterial communities in whole mosquito samples in function of A: *Wolbachia* infection (relative abundance – HCA) and *Wolbachia* density (qPCR) B: Mosquito species C: Mosquito strain D: Field and laboratory strains.

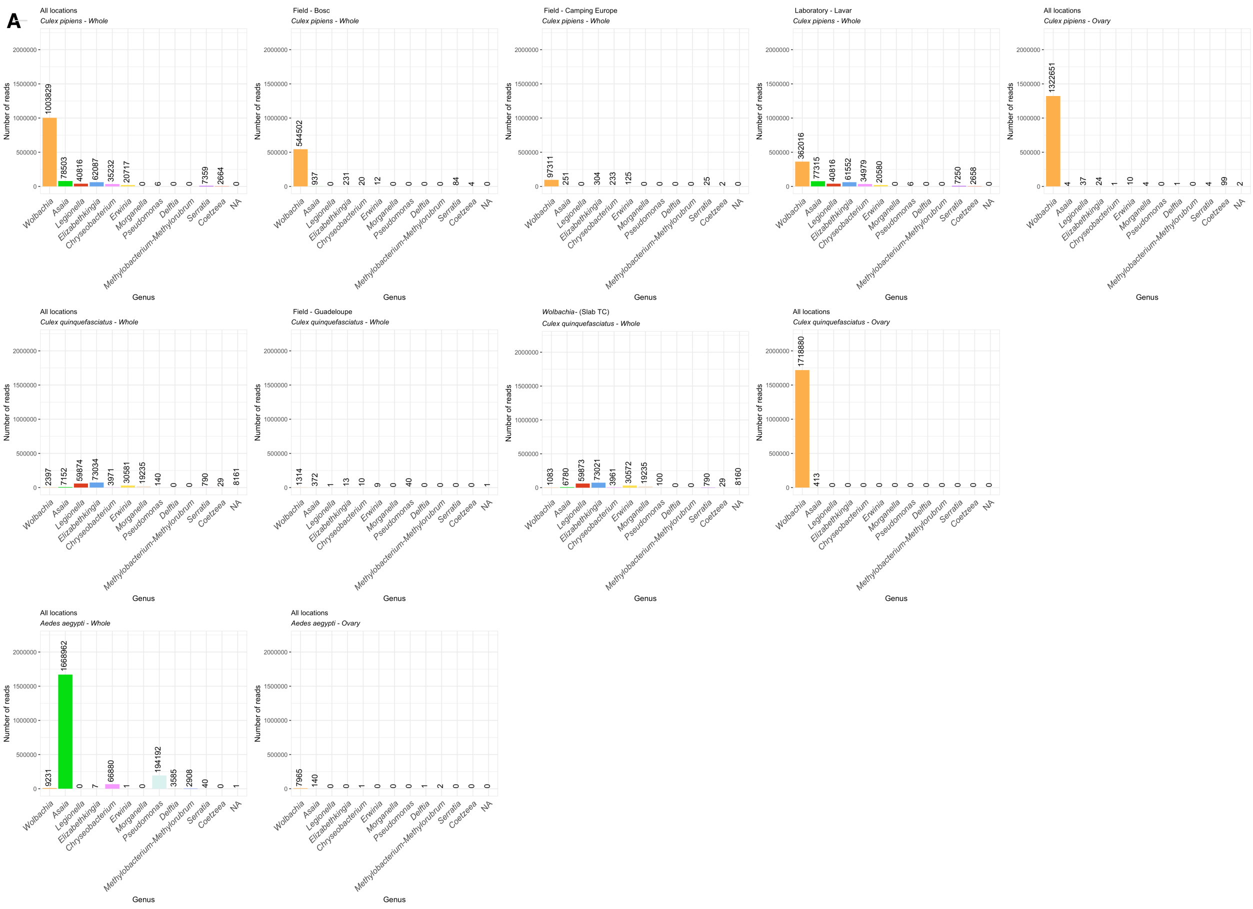

A
All locations
Culex pipiens - Whole
1003829 78503 40816 62087 35232 20717 0 6 0 0 7359 2664 0
Field - Bosc
Culex pipiens - Whole
544502 937 0 231 20 12 0 0 0 0 84 4 0
Field - Camping Europe
Culex pipiens - Whole
97311 251 0 304 233 125 0 0 0 0 25 2 0
Laboratory - Lavar
Culex pipiens - Whole
362016 77315 40816 61552 34979 20580 0 6 0 0 7250 2658 0
All locations
Culex pipiens - Ovary
1322651 4 37 24 1 10 4 0 1 0 4 99 2
All locations
Culex quinquefasciatus - Whole
2397 7152 59874 73034 3971 30581 19235 140 0 0 790 29 8161
Field - Guadeloupe
Culex quinquefasciatus - Whole
1314 372 1 13 10 9 0 40 0 0 0 0 1
Wolbachia- (Slab TC)
Culex quinquefasciatus - Whole
1083 6780 59873 73021 3961 30572 19235 100 0 0 790 29 8160
All locations
Culex quinquefasciatus - Ovary
1718880 413 0 0 0 0 0 0 0 0 0 0 0
All locations
Aedes aegypti - Whole
9231 1668962 0 7 66880 1 0 194192 3585 2908 40 0 1
All locations
Aedes aegypti - Ovary
7965 140 0 0 1 0 0 0 1 2 0 0 0
Number of reads
0 500000 1000000 1500000 2000000
Wolbachia Asaia Legionella Elizabethkingia Chryseobacterium Erwinia Morganella Pseudomonas Delftia Methylobacterium-Methylorubrum Serratia Coetzeea NA
Genus

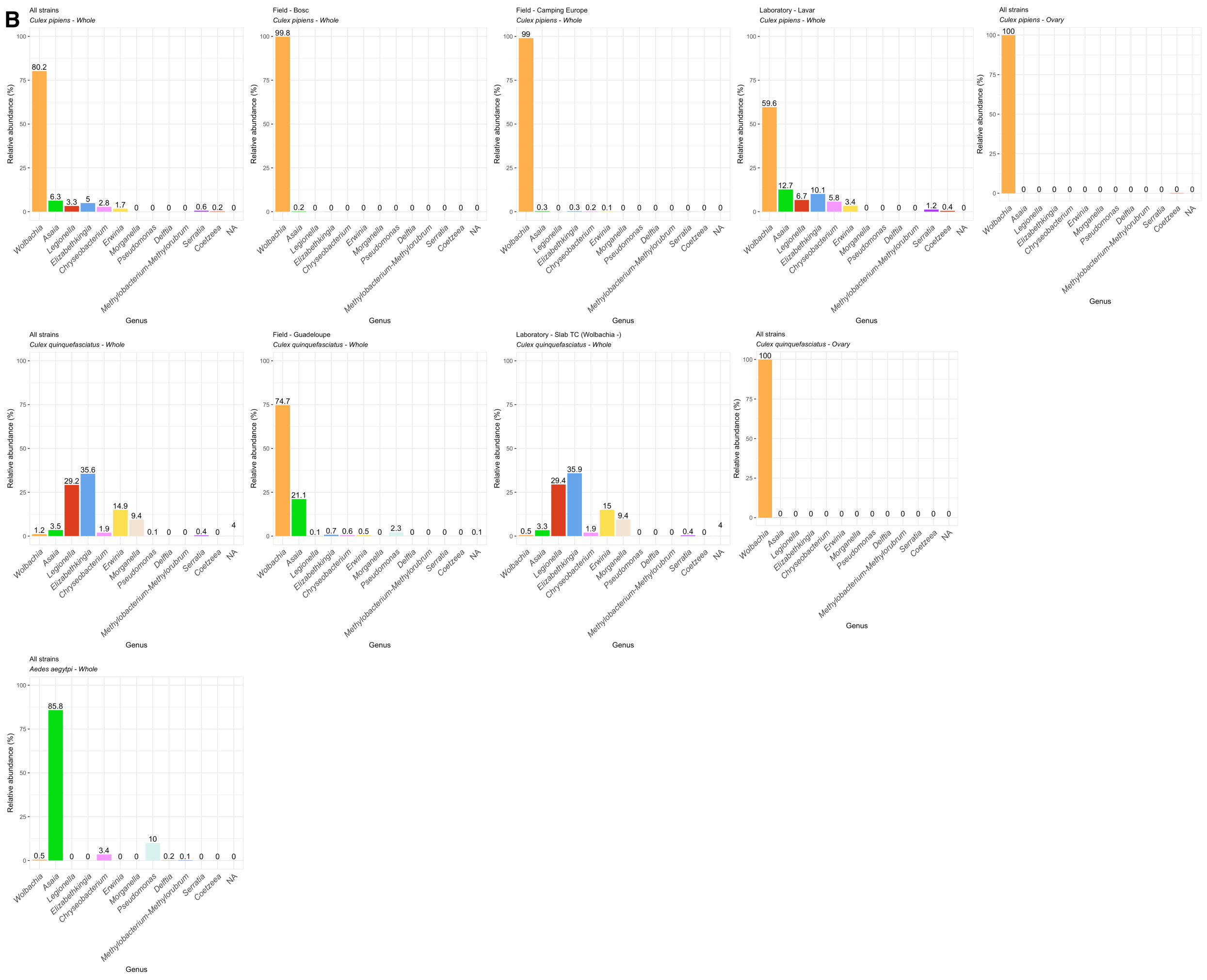


**Supplementary Figure 8**: Panel A - Number of reads per genus for the 113 samples retained after MED analysis, decontam and the removing of samples with less than 1000 reads (with the exception of NP20, NP36, NP29, NP30, NP34 as detailed in Supplementary Note 1) for the different species *Culex pipiens*, *Culex quinquefasciatus*, *Aedes aegypti* for all vs. each separate strain, whole and ovary samples (each subpanel) . Panel B - Rounded relative abundance (at 1 decimal) of reads per genus for the different species *Culex pipiens*, *Culex quinquefasciatus*, *Aedes aegypti* for all vs. each separate strain, whole and ovary samples (each subpanel). Of note, one of the two *Aedes* ovary samples showed very low number of reads (140), indicating some misleading percentage numbers for this group of samples (as shown by rarefaction curves that did not reach a plateau and incomplete sequencing depth) and was therefore not shown.

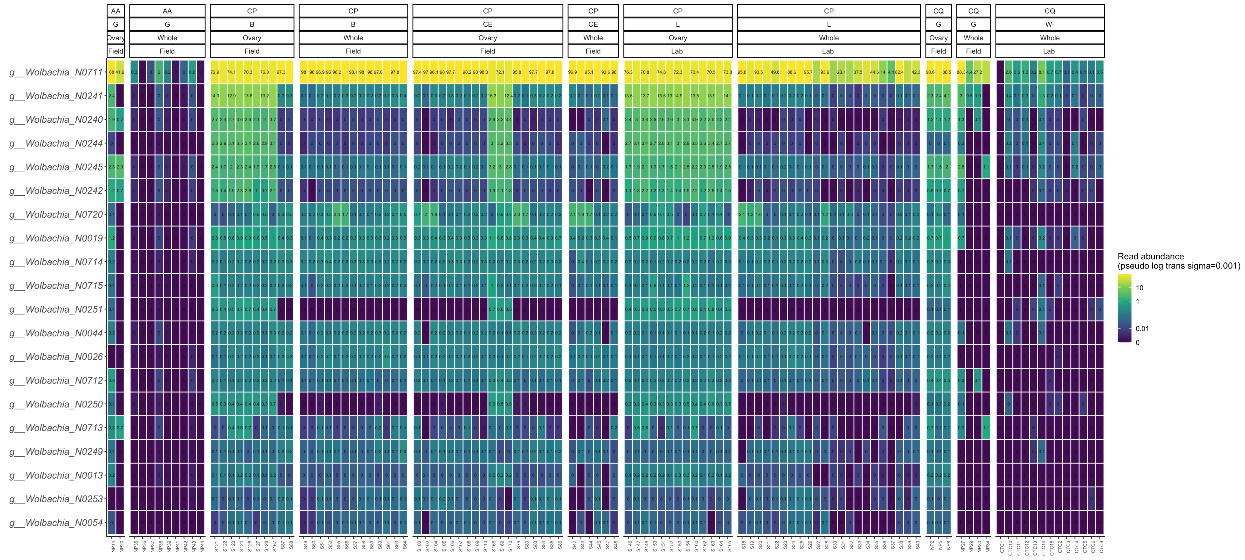


**Supplementary Figure 9**: Heatmap of unsupervised oligotypes assigned to *Wolbachia* in samples (pseudo log transformation with sigma=0.001). AA=*Aedes aegypti*, CP=*Culex pipiens*, CQ=*Culex quinquefasciatus*, G=Guadeloupe, B=Bosc, CE= Camping Europe, L=Lavar and W-=Slab TC Wolbachia-.

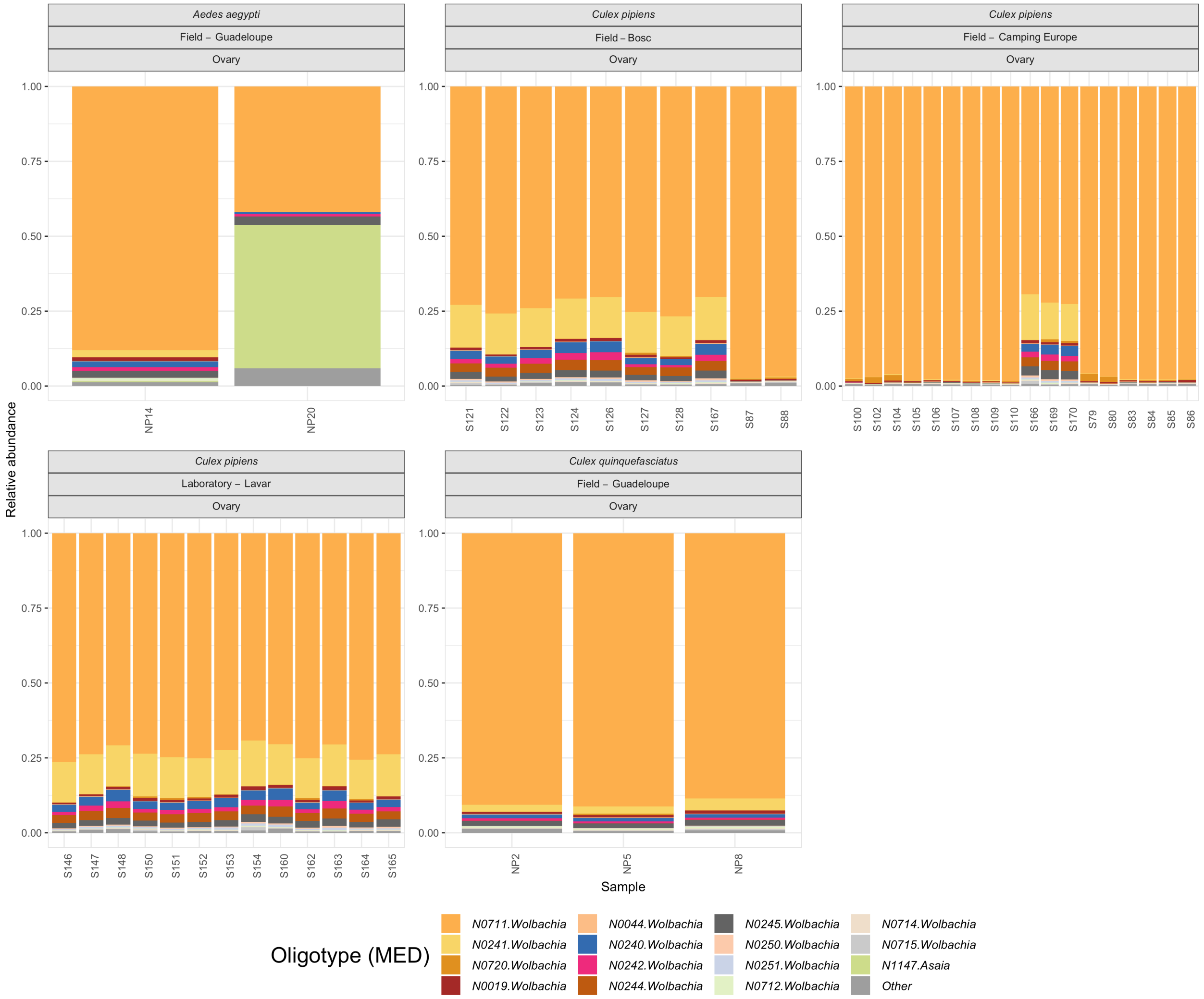


**Supplementary Figure 10**: The 15 most abundant unsupervised oligotypes in the ovaries of *Culex* spp and *Aedes aegypti* individuals. Each color represents an unsupervised oligotype with his taxonomic assignment at the genus level. "Other" groups indicate unsupervised oligotypes that are not included in the 15 most abundant oligotypes.

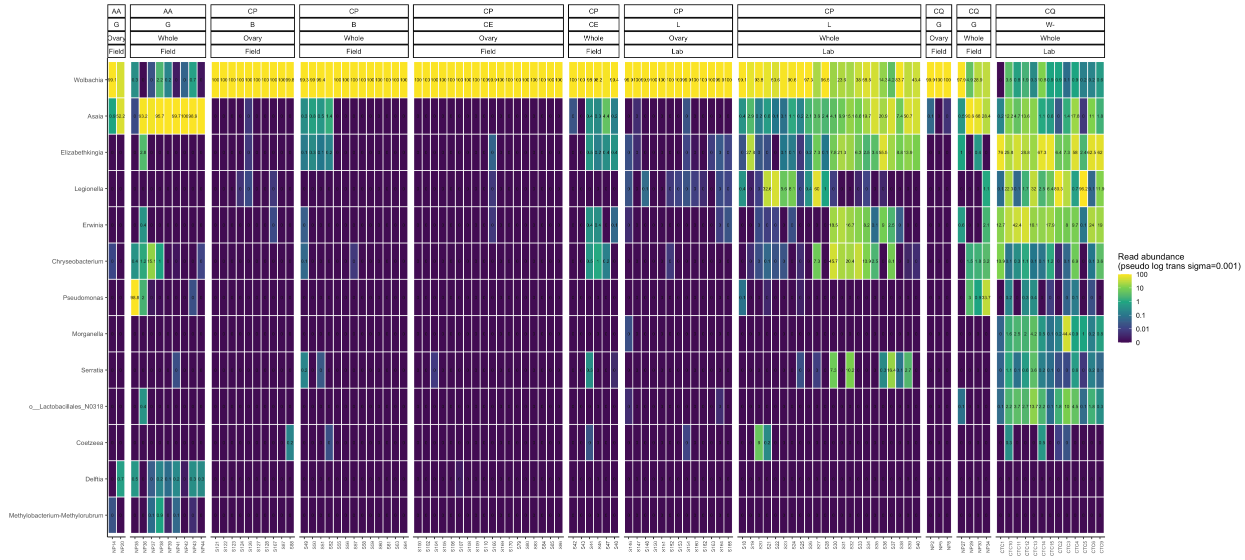


**Supplementary Figure 11**: Heatmap of unsupervised oligotypes for each genus in the dataset (pseudo log transformation with sigma=0.001). AA=*Aedes aegypti,* CP=*Culex pipiens*, CQ=*Culex quinquefasciatus*, G=Guadeloupe, B=Bosc, CE= Camping Europe, L=Lavar and W- =Slab TC Wolbachia-.

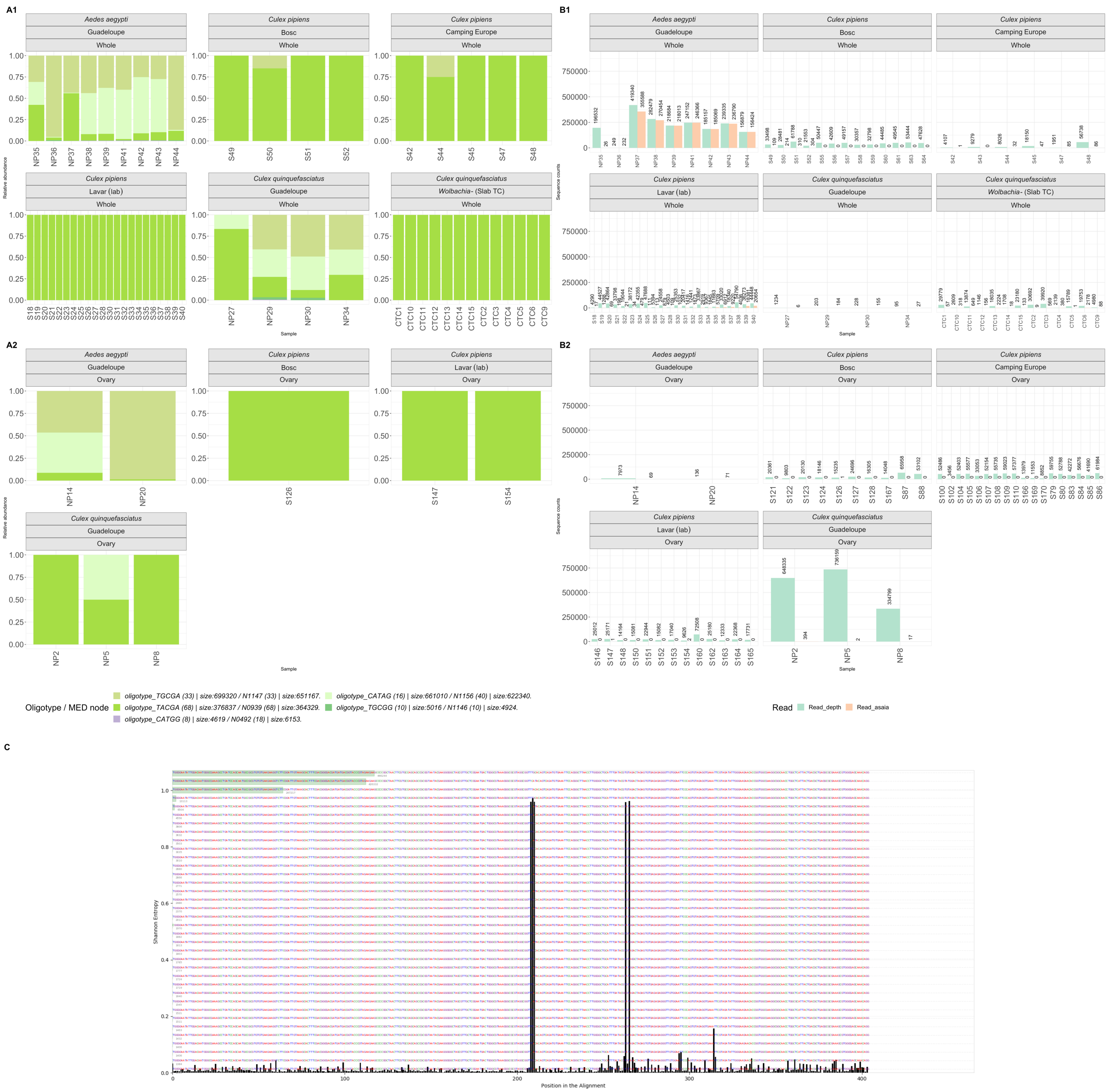


**Supplementary Figure 12**: A. Relative abundance of the *Asaia* supervised oligotypes in whole mosquito samples (A1) and in ovary samples (A2). Each color represents a supervised oligotype, its frequency across the different samples and size (number of reads) followed by the corresponding unsupervised oligotype post-MED analysis with its respective frequency and size. B. Number of the total reads (green) and number of *Asaia* reads (orange) in whole samples (B1) and in ovary samples (B2). C. Shannon Entropy values for each nucleotide position for the *Asaia* reads.

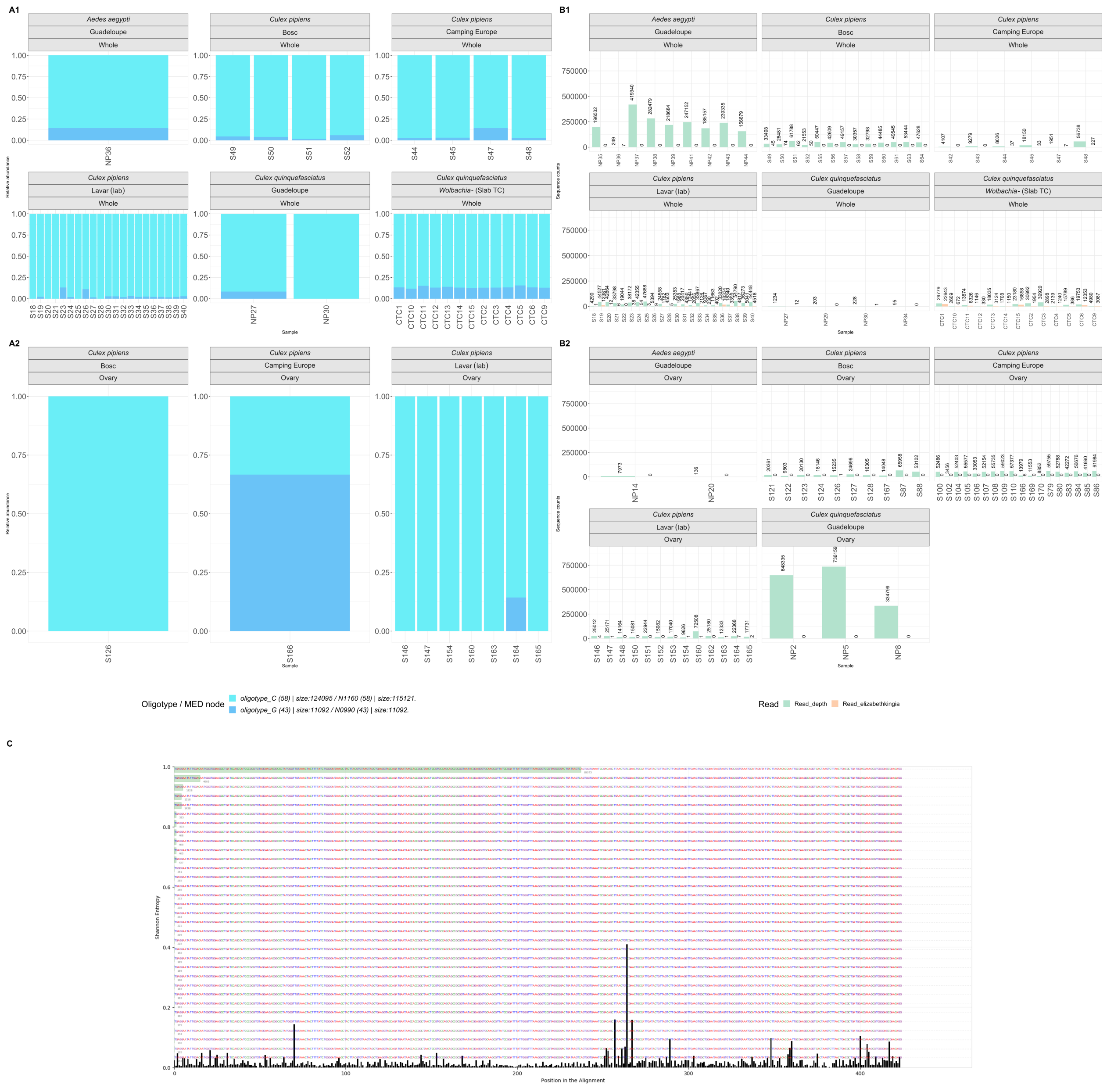


**Supplementary Figure 13**: A. Relative abundance of the *Elizabethkingia* supervised oligotypes in whole mosquito samples (A1) and in ovary samples (A2). Each color represents a supervised oligotype, its frequency across the different samples and size (number of reads) followed by the corresponding unsupervised oligotype post-MED analysis with its respective frequency and size. B. Number of the total reads (green) and number of *Elizabethkingia* reads (orange) in whole samples (B1) and in ovary samples (B2). C. Shannon Entropy values for each nucleotide position for the *Elizabethkingia* reads.

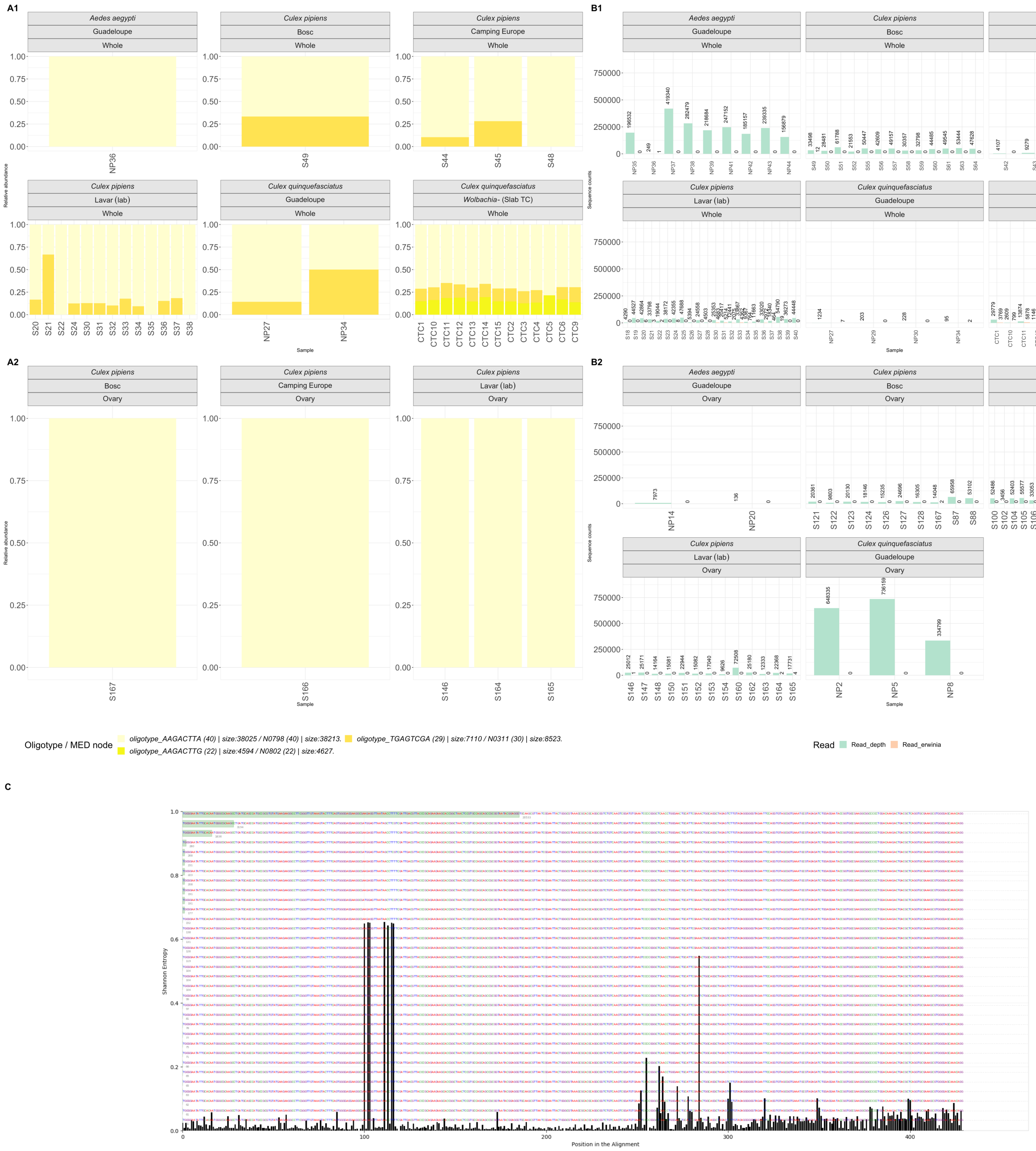

**Supplementary Figure 14:** A. Relative abundance of the *Erwinia* supervised oligotypes in whole mosquito samples (A1) and in ovary samples (A2). Each color represents a supervised oligotype, its frequency across the different samples and size (number of reads) followed by the corresponding unsupervised oligotype post-MED analysis with its respective frequency and size. B. Number of the total reads (green) and number of *Erwinia* reads (orange) in whole samples (B1) and in ovary samples (B2). C. Shannon Entropy values for each nucleotide position for the *Erwinia* reads.

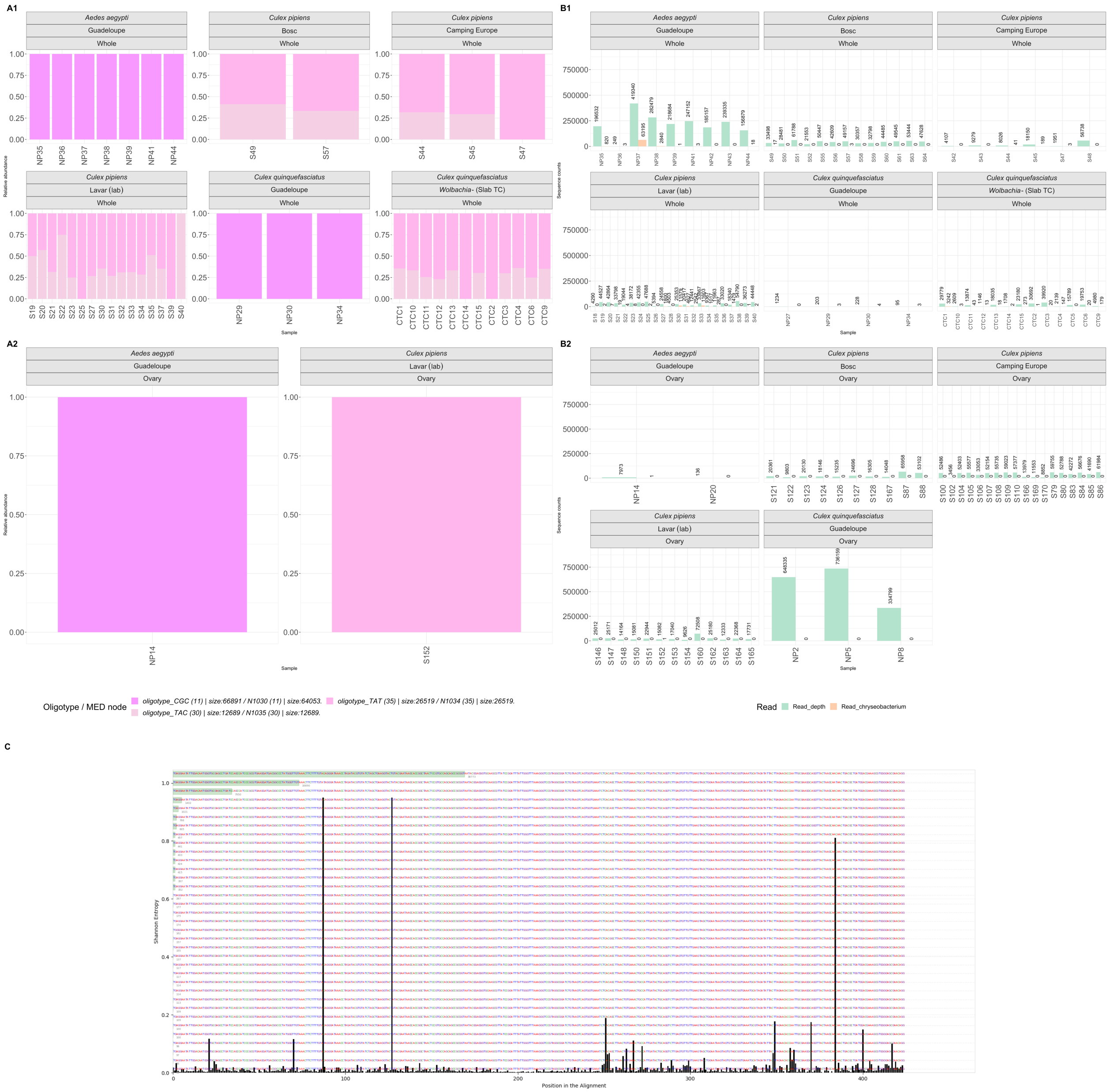


**Supplementary Figure 15 :** A. Relative abundance of the *Chryseobacterium* supervised oligotypes in whole mosquito samples (A1) and in ovary samples (A2). Each color represents a supervised oligotype, its frequency across the different samples and size (number of reads) followed by the corresponding unsupervised oligotype post-MED analysis with its respective frequency and size. B. Number of the total reads (green) and number of *Chryseobacterium* reads (orange) in whole samples (B1) and in ovary samples (B2). C. Shannon Entropy values for each nucleotide position for the *Chryseobacterium* reads.

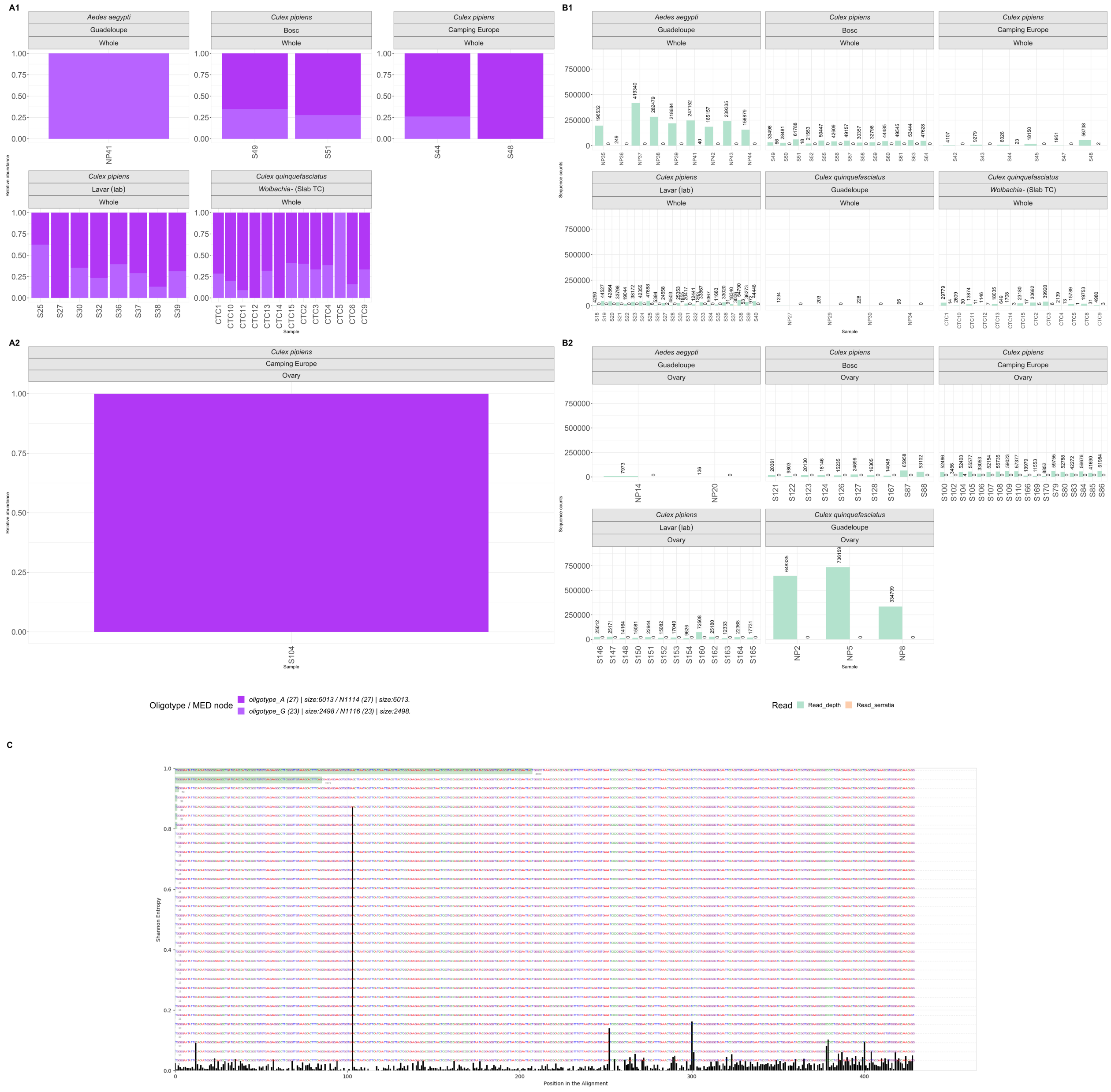


**Supplementary Figure 16:** A. Relative abundance of the *Serratia* supervised oligotypes in whole mosquito samples (A1) and in ovary samples (A2). Each color represents a supervised oligotype, its frequency across the different samples and size (number of reads) followed by the corresponding unsupervised oligotype post-MED analysis with its respective frequency and size. B. Number of the total reads (green) and number of *Serratia* reads (orange) in whole samples (B1) and in ovary samples (B2). C. Shannon Entropy values for each nucleotide position for the *Serratia* reads.

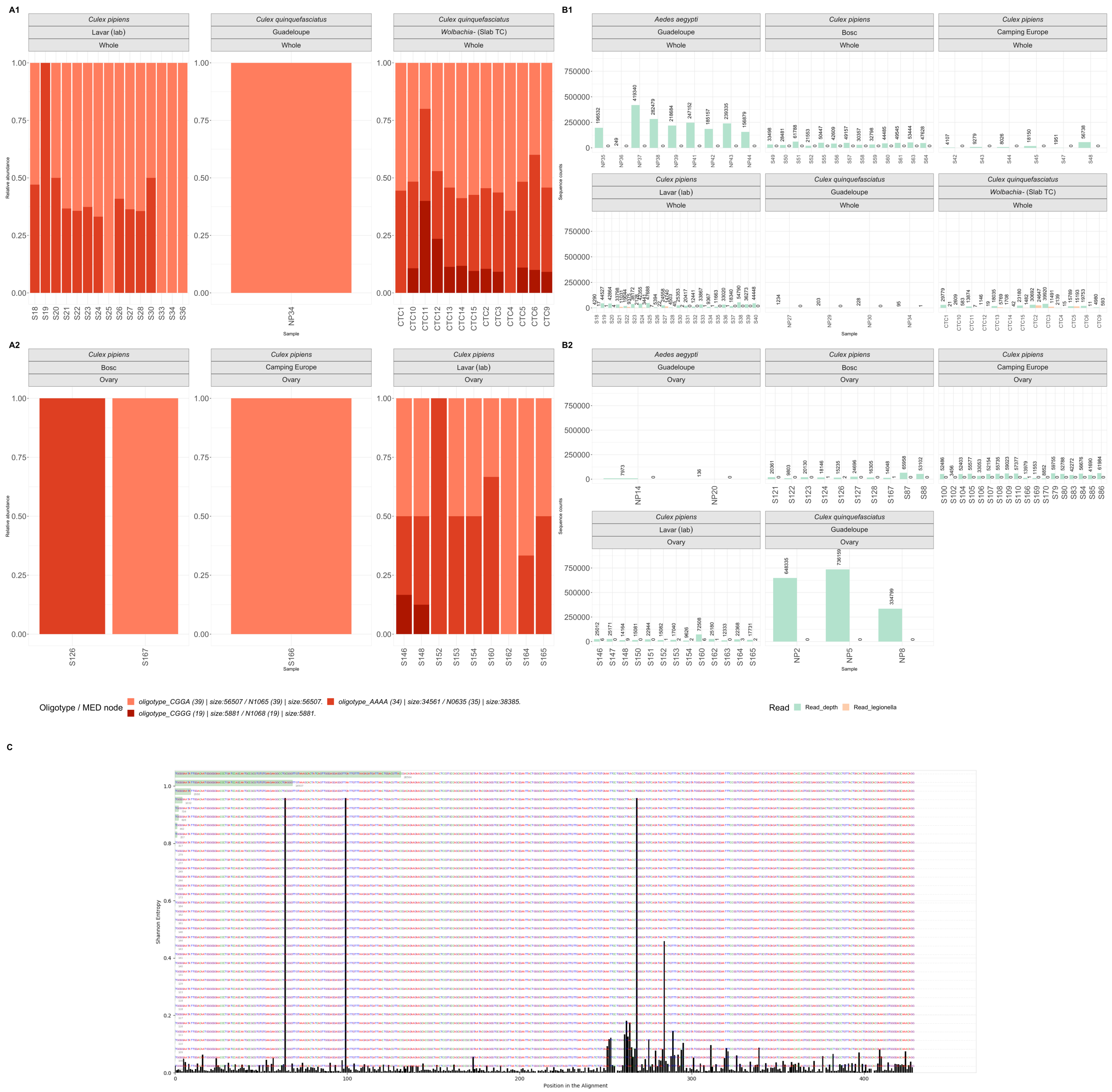


**Supplementary Figure 17**: A. Relative abundance of the *Legionella* supervised oligotypes in whole mosquito samples (A1) and in ovary samples (A2). Each color represents a supervised oligotype, its frequency across the different samples and size (number of reads) followed by the corresponding unsupervised oligotype post-MED analysis with its respective frequency and size. B. Number of the total reads (green) and number of *Legionella* reads (orange) in whole samples (B1) and in ovary samples (B2). C. Shannon Entropy values for each nucleotide position for the *Legionella* reads.

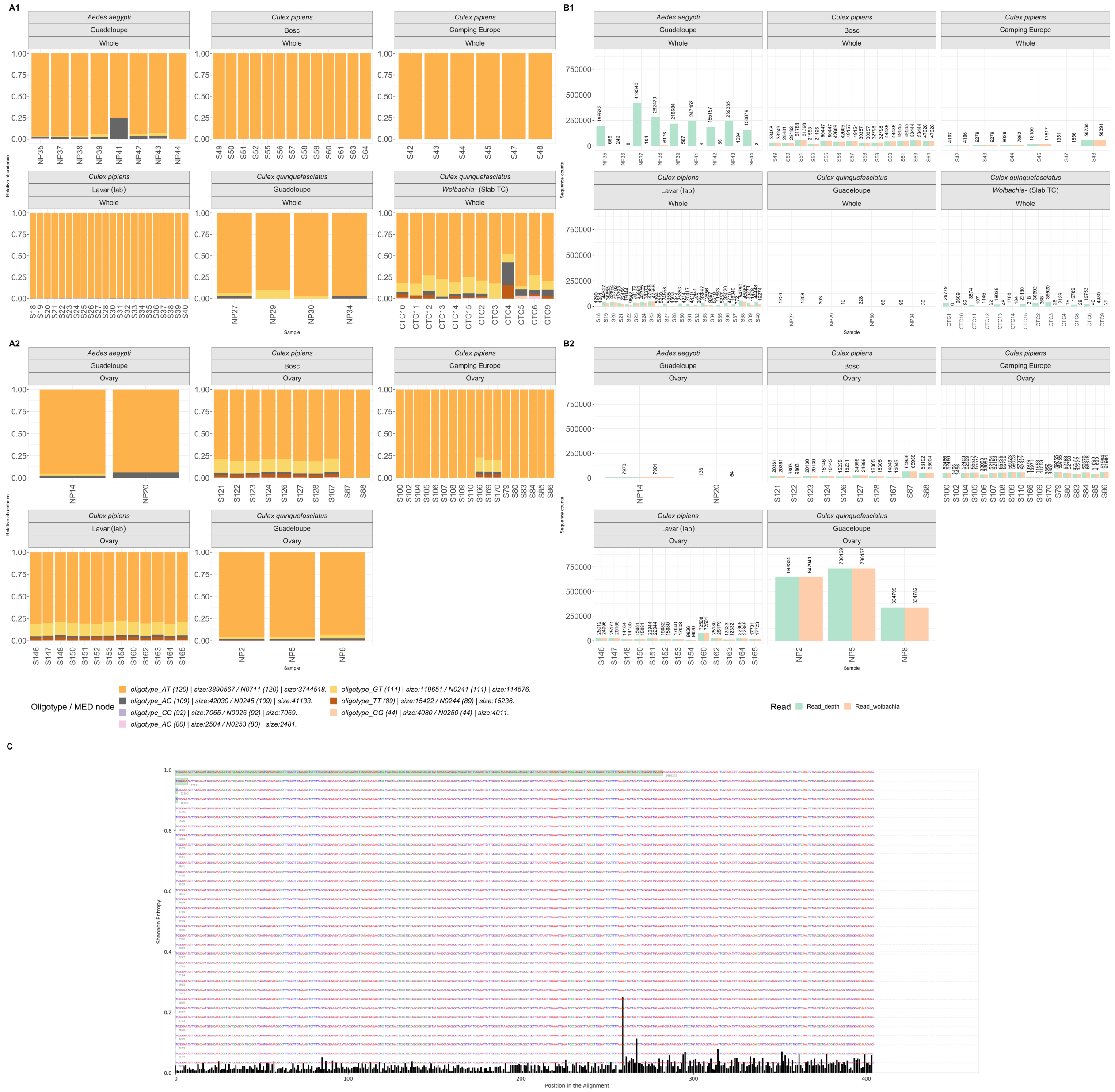


**Supplementary Figure 18**: A. Relative abundance of the *Wolbachia* supervised oligotypes in whole mosquito samples (A1) and in ovary samples (A2). Each color represents a supervised oligotype, its frequency across the different samples and size (number of reads) followed by the corresponding unsupervised oligotype post-MED analysis with its respective frequency and size. B. Number of the total reads (green) and number of *Wolbachia* reads (orange) in whole samples (B1) and in ovary samples (B2). C. Shannon Entropy values for each nucleotide position for the *Wolbachia* reads.

| Oligotype | Fragments (30bp) |
|---|---|
| AT | TGCTTTTAAAACTATTAATCTAGAGATTGA |
| GT | TGCTTTTAAAGCTATTAATCTAGAGATTGA |
| AG | TGCTTTTAAAACTATTAAGCTAGAGATTGA |
| TT | TGCTTTTAAATCTATTAATCTAGAGATTGA |
| CC | TATTGGGCGTAAGGGCGCGTAGGCTGGTTA |
| GG | TGCTTTTAAAGCTATTAAGCTAGAGATTGA |
| AC | TGCTTTTAAAACTATTAACCTAGAGATTGA |

**Supplementary Figure 19**: Fragments extracted from each *Wolbachia* supervised oligotype used to find hits in the seven metagenomes from *Culex pipiens* individuals. Colored letters correspond to high entropy positions.

| | AT | GT | AG | TT | CC | GG | AC |
|---|---|---|---|---|---|---|---|
| Culex O03 | 172 | 0 | 1 | 0 | 0 | 0 | 0 |
| Culex O07 | 146 | 0 | 0 | 0 | 0 | 0 | 0 |
| Culex O11 | 333 | 1 | 1 | 0 | 0 | 0 | 0 |
| Culex O12 | 96 | 0 | 0 | 0 | 0 | 0 | 0 |
| Pipiens_MGx_Istanbul | 298 | 0 | 0 | 0 | 0 | 0 | 0 |
| Pipiens_MGx_Tunis | 347 | 0 | 0 | 0 | 1 | 0 | 0 |
| Pipiens_MGx_Harash | 319 | 0 | 0 | 0 | 0 | 0 | 0 |

**Supplementary Figure 20**: Hits of *Wolbachia* supervised oligotype fragments (30bp) in the four quality-filtered *Culex pipiens* ovary metagenomes from Reveillaud et al., 2019[1] (European Nucleotide Archive ENA: ERS2407346, Culex O03; ERS2407347, Culex O07; ERS2407348, Culex O11; ERS2407349, Culex O12) and the three isofemale lines *C. pipiens* egg-rafts from North Africa from Bonneau et al., 2018[2] (SRR5810516, Pipiens_MGx_Istanbul; SRR5810518, Pipiens_MGx_Tunis; SRR5810517 Pipiens_MGx_Harash).

[1] Reveillaud et al., "The *Wolbachia* mobilome in *Culex pipiens* includes a putative plasmid." *Nat Commun*. 2019;10(1):1051
[2] Bonneau et al., "*Culex pipiens* crossing type diversity is governed by an amplified and polymorphic operon of *Wolbachia*." *Nat Commun*. 2018;9(1):319.
